# The origin of taxonomically restricted genes in yeast

**DOI:** 10.64898/2026.08.24.746667

**Authors:** Aaron Wacholder, Anne-Ruxandra Carvunis

## Abstract

The evolutionary origins of taxonomically restricted “orphan” genes (TRGs), which lack known homologs outside of a specific taxon, have fueled a decades-long controversy. On the one hand, thousands of TRGs are proposed to have originated *de novo* from previously non-genic DNA. These include an expansive repertoire of newly discovered microproteins that were missed from genome annotations yet can mediate important phenotypes. On the other hand, critics contend that many TRGs are not novel gene creations, but rather the products of extreme sequence divergence leading to homology detection failure. Here, we resolve this controversy by systematically dissecting the origins of TRGs across the *Saccharomyces* taxon, using a sensitive profile-profile alignment approach to identify challenging homologs. Our results reconcile the two competing models by revealing a temporal shift in the mechanisms underlying TRG formation. We demonstrate that most species-specific TRGs are genuine *de novo* gene births whereas most of the TRGs that are conserved across the *Saccharomyces* genus derive from highly diverged ancestral genes whose homology is no longer detectable using common methods. These findings suggest that, in *Saccharomyces*, a high rate of *de novo* birth events is balanced by evolutionary attrition with little to no survivors after a few million years. Therefore, nearly all *de novo* genes appear destined to vanish, with little contribution to the stable genetic repertoire over deep evolutionary time, despite providing important contributions to species-specific physiology and adaptation in the present time.

## Introduction

One of the most surprising results from early genome sequencing projects was the prevalence of “orphan” protein-coding genes with no known homologs in other organisms.^1^ Even as the number of sequenced genomes has exploded, we continue to find that each genome contains many “taxonomically-restricted genes” (TRGs) with no identifiable homologs outside a narrow taxonomic range.^2,3^ All genomes appear to contain genes restricted to the species, genus, family, and each further taxonomic rank. The recent discovery that eukaryotic genomes encode hundreds to thousands of microproteins, mostly not annotated in gene databases and generally lacking known homologs outside closely related species, expands the list of TRGs still further.^4–6^

Over the past two decades, controversy has emerged over how to interpret the ubiquity of TRGs. On the one hand, the prevalence of TRGs may suggest a major role for *de novo* gene birth in evolution.^7–9^ A gene born *de novo* within a given taxon will have no homologs outside the taxon, so frequent *de novo* birth can explain why many genes appear taxonomically restricted. On the other hand, there is an alternative explanation for TRGs: a gene could have homologs outside the taxon that are missed by homology detection algorithms due to extreme sequence divergence, a phenomenon termed homology detection failure.^10–13^ Alternative statistical models suggest different levels of concern for homology detection failure.^14^ Weisman et al. 2020 found that most TRGs can be explained by homology detection failure due to their typically short lengths and rapid evolutionary rates.^12^ Yet, Vakirlis et al. 2020, using an alternative approach, predicted that a majority of TRGs are not explained by homology detection failure.^15^ The assumption that most TRGs are of *de novo* origin underlies evolutionary modeling of the trajectory of *de novo* genes from birth to development into mature genes.^7,16–19^ Without a resolution to this controversy, the contribution of *de novo* gene birth to new gene formation is unclear.^20^

The recent availability of hundreds of genomes from the budding yeast subphylum^21,22^ provides an opportunity to assess the origin of TRGs through direct identification of outgroup homologs. Here, focusing on genes that initially appear restricted to the *Saccharomyces* genus in a BLAST search, we combine powerful profile-profile alignment methods with systematic use of synteny to achieve more sensitive homology detection than previously possible and thus resolve the evolutionary history of a large majority of *Saccharomyces*-restricted genes.

We find that the mechanisms leading to taxonomic restriction differ greatly by taxonomic rank. TRGs restricted to taxa below the *Saccharomyces* clade are overwhelmingly of recent *de novo* origin. However, TRGs conserved across the *Saccharomyces* genus are mostly explained by homology detection failure. These findings can be explained under a model where *de novo* gene birth is common, creating an abundance of species-specific genes, but these genes almost never persist over more than a few million years to become TRGs of higher taxonomic rank.

## Results

### Identifying Saccharomyces-restricted genes in Saccharomyces cerevisiae

To better understand the origins of TRGs, we first identified the TRGs in *S. cerevisiae* restricted to the *Saccharomyces* genus (Figure 1A-B, Supplementary Figure 1A). We consider a gene to be a *Saccharomyces*-restricted gene (SRG) if we cannot exclude the possibility that it is unique to the *Saccharomyces* genus using BLAST^23^ searches. For each annotated *S. cerevisiae* gene, we used BLASTP and TBLASTN to search for homologs against 1,146 yeast genomes assembled by Opulente et al. 2024.^21^ A gene was inferred to have an extrageneric homolog if it had 1) a BLASTP or TBLASTN match with e-value < 10^−4^ with at least three species outside the genus, or 2) e-value < 10^−6^ with at least one such species or 3) an e-value < 10^−6^ to another gene in *S. cerevisiae* that met one of the first two criteria.^24^ Using scrambled sequences of annotated genes as controls, we find that these e-value thresholds provide a false discovery rate < 0.2% (Supplementary Figure 1B). We identify 310 genes in *S. cerevisiae* that meet our definition of SRG (Figure 1B, Supplementary Figure 1C-F, Supplementary Table 1).

**Figure 1:**
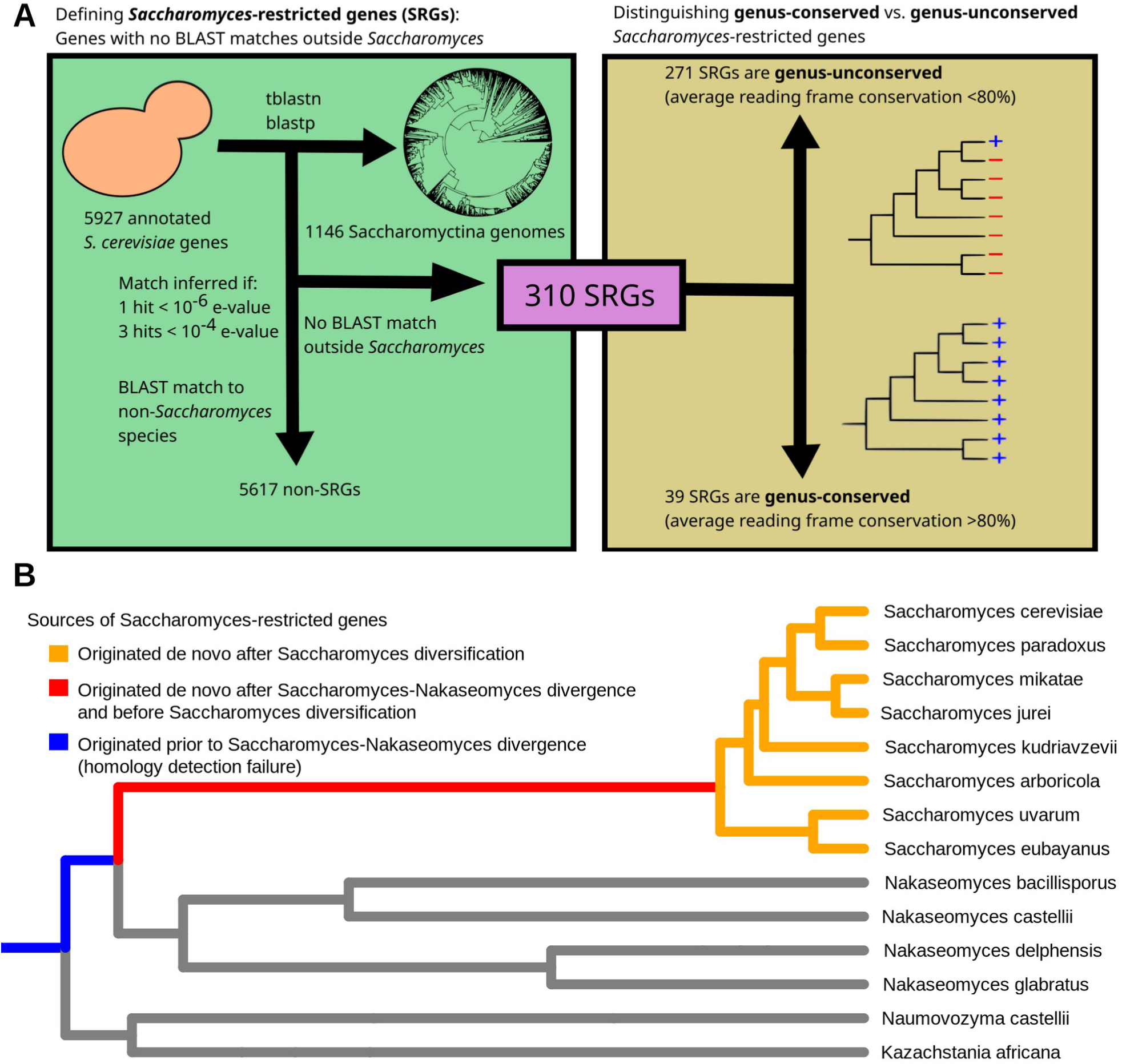
Identification and classification of Saccharomyces-restricted genes. A) Flowchart describing the identification of 310 Saccharomyces-restricted genes (SRGs) and their partitioning into genus-conserved and genus-unconserved classes based on reading frame conservation within Saccharomyces. B) Phylogenetic tree of the Saccharomyces genus and its most closely related genera, indicating the potential origins of SRGs. SRGs that emerged de novo after Saccharomyces-Nakaseomyces divergence will genuinely lack any homologs outside Saccharomyces, while those originating before will generally have homologs outside the genus that were not detected by BLAST.

### A minority of SRGs are conserved within the ***Saccharomyces*** genus

SRGs can be divided into those restricted at the genus level but not below—i.e. SRGs conserved across *Saccharomyces*—and those also restricted to lower taxonomic levels. We identify genus-conserved SRGs using reading frame conservation (RFC), a measure developed by Kellis et al. 2003^25^ to identify open reading frames (ORFs) with conserved reading frames between *S. cerevisiae* and its congeneric species. Across all *S. cerevisiae* ORFs, average RFC between an ORF and its orthologous loci in other *Saccharomyces* species shows a bimodal distribution, with ORFs conserved across the genus (mean RFC > 80%) cleanly distinguished from those that lack genus-level conservation (mean RFC < 60%).^4^ We similarly observe a bimodal distribution of mean RFC among SRGs (Figure 2A). We therefore consider an SRG to be conserved within the genus if it has mean RFC > 80% among its orthologous loci in other *Saccharomyces* species, while SRGs with mean RFC < 80% are not conserved across the genus. By this definition, 5110 of 5284 (96.7%) of annotated single-exon yeast non-SRGs are conserved within *Saccharomyces*. In contrast, among the 310 SRGs, only 39 (12.6%) are conserved within *Saccharomyces* while most (271) are restricted at lower taxonomic ranks. This reflects a longstanding observation that species-specific TRGs are disproportionately numerous.^3^

**Figure 2:**
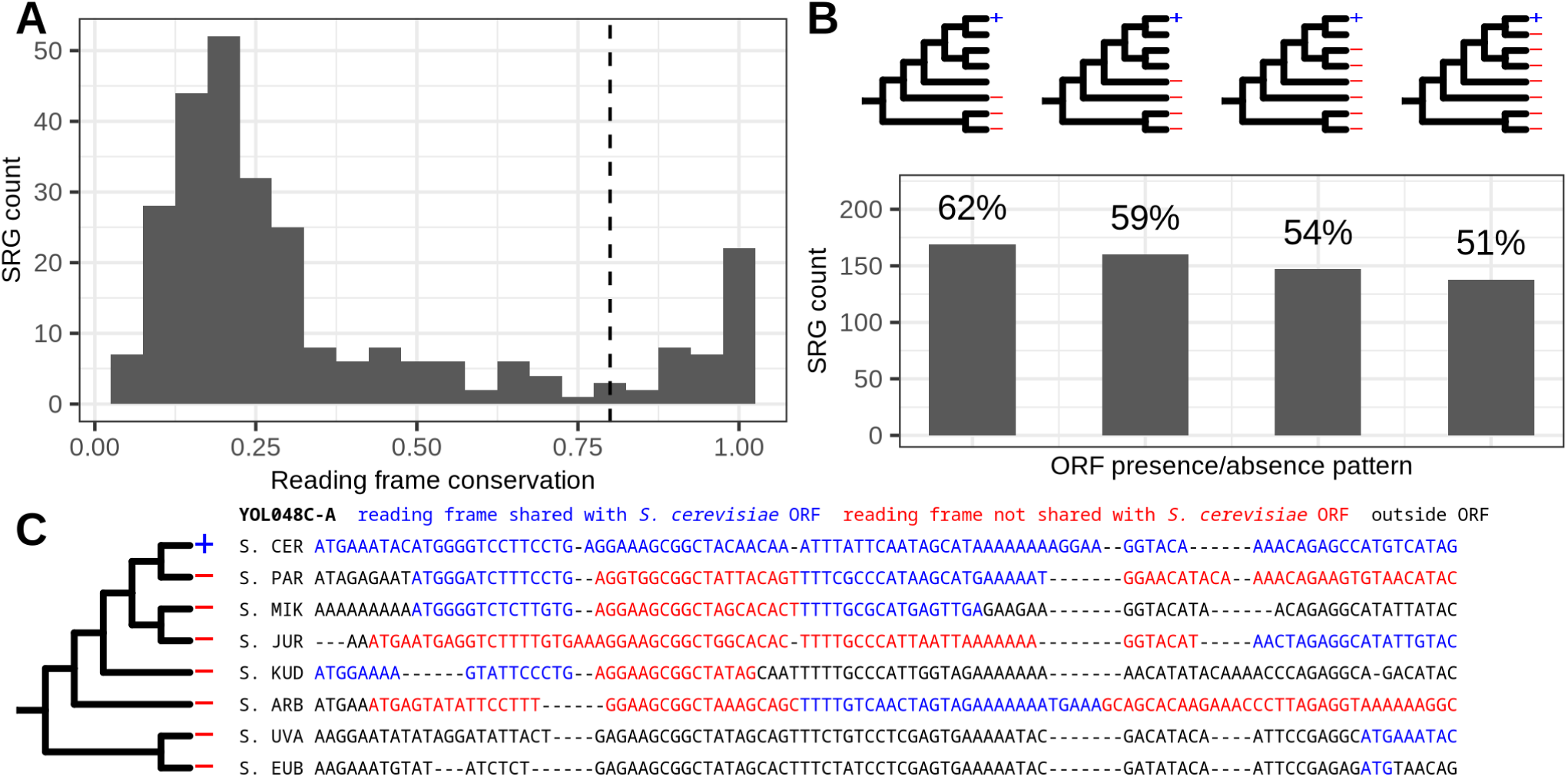
Most Saccharomyces-restricted genes that lack genus-level conservation are of de novo origin. A) Distribution of mean reading frame conservation values among SRGs that have orthologous DNA sequence in at least four other Saccharomyces species (N=277). The reading frame conservation indicated is an average of the S. cerevisiae gene against all its orthologous DNA sequences in Saccharomyces. Reading frame conservation above 0.8, indicated by a dashed line, is taken to indicate conservation within Saccharomyces and delineate a class of 39 genus-conserved SRGs. B) The number and percentage (indicated above bar) of genus-unconserved SRGs matching specified patterns (top) of presence/absence in Saccharomyces. A blue + sign on the tree indicates the gene must be present in that species for the gene to match the pattern and a red – sign indicates the gene must be absent in that species. The gene is considered present in a species if an ORF exists at the orthologous locus with reading frame conservation > 0.8 with the S. cerevisiae gene. All patterns indicate that de novo origin is supported by parsimony, with patterns further to the right providing increasingly stronger evidence. C) Example alignment of a gene, YOL048C-A, with an ORF presence/absence pattern providing strong support for de novo origin. Though the orthologous region is present in all Saccharomyces species with identifiable homology at the nucleotide level, frameshifts, missing starts, and early stops indicate that the S. cerevisiae reading frame is poorly conserved in all other Saccharomyces species. The ORF represented in each species is that with the highest reading frame conservation with the S. cerevisiae ORF.

### Most SRGs that lack genus-level conservation are of recent *de novo* origin

We next assessed the evidence for *de novo* origin among the 271 SRGs that are not conserved within the genus. Confident inference of *de novo* origin requires identifying the orthologous DNA sequence of the *S. cerevisiae* SRG locus in outgroup lineages and demonstrating that the ORF is absent from the orthologous locus, such that an ancestral noncoding state can be inferred. The more such outgroup lineages can be identified, the higher confidence can be obtained that the ancestral state was non-genic rather than an ancestral coding state that subsequently experienced losses in some lineages. In 169 of 271 (62%) SRGs the ORF is absent at the orthologous locus in *S. uvarum*, *S. eubayanus*, and *S. arboricola*, such that parsimony supports *de novo* origin (Figure 2B). For 138 of 271 SRGs (51%) the ORF is absent in every species of *Saccharomyces* other than *S. cerevisiae*, providing high confidence that the ORF is a species-specific gene of recent *de novo* origin (Figure 2B-C, Supplementary Table 1). We conclude that *de novo* gene birth is the most important contributor to SRGs that lack genus-level conservation.

To determine whether any of the 39 SRGs that are conserved within *Saccharomyces* are of *de novo* origin would require identifying orthologous regions for these SRGs in *Nakaseomyces*, the most closely related genus to *Saccharomyces* in the Opulente et al. 2024 genome dataset.^21^ However, *Nakaseomyces* is too distant from *Saccharomyces* for nucleotide homology to be generally identifiable; there are no BLASTN hits to any of the 39 genus-conserved SRGs at a 10^−4^ e-value threshold. Further understanding the origin of genus-conserved SRGs therefore requires alternative approaches.

### Sensitive homology detection identifies vertically descended SRGs with preserved synteny

We next considered the extent to which SRGs could be derived by vertical descent. To improve homology detection for vertically descend SRGs, we made use of two ideas. First, we used synteny to attempt to detect homologs of SRGs within the region we expect to find them, improving power relative to a whole-genome search. Second, we used profile-profile alignments to test homology between the multiple sequence alignments (MSAs) of SRGs against candidate gene MSAs in other yeast genera, improving power relative to pairwise alignments.^26^ Combining these ideas, we developed an automated method to search syntenic regions of post-whole genome duplication (WGD) genera for homologs of SRGs using profile-profile alignment from hh-suite (See Methods, Supplementary Figure 2). To control false discovery rate, we also searched for homologs for each SRG in the syntenic regions of other SRGs; we have no reason to expect genuine homology in these alignments and so any hit is expected to be a false positive.

Our automated pipeline detected syntenic matches between 19 of 39 (49%) genus-conserved SRGs to a total of 44 homology groups outside *Saccharomyces* at a 5% e-value threshold (FDR < 5%, Figure 3A). In contrast, among SRGs that lack genus-level conservation, (N=109; excluding those that overlap any other annotated ORF or functional RNA), none had any matches at this threshold (Figure 3A). Thus, around half (49%) of SRGs conserved within *Saccharomyces* have apparent syntenic homologs outside *Saccharomyces,* while SRGs that are not conserved within the genus show no evidence of extrageneric syntenic homologs.

**Figure 3:**
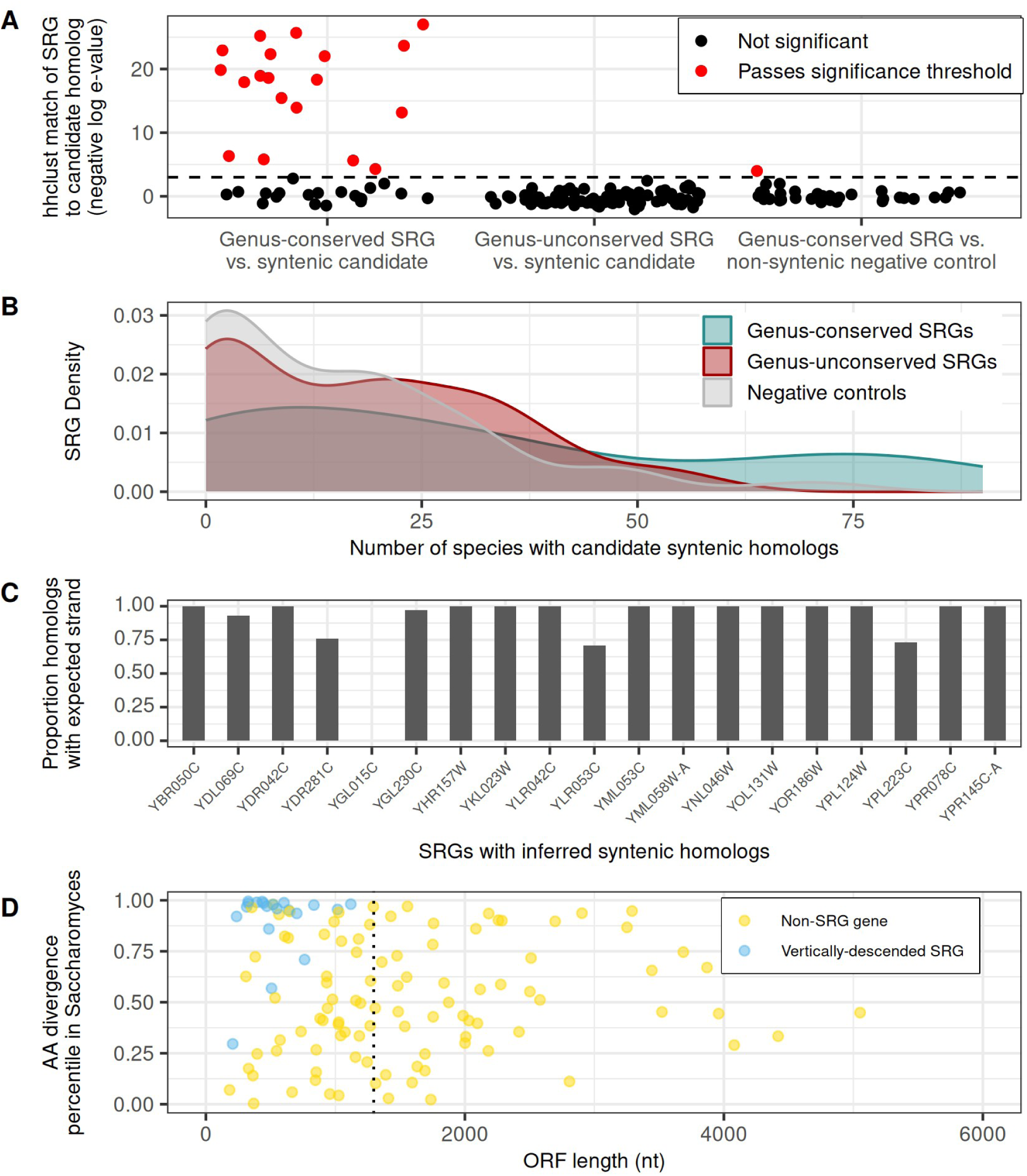
Profile-profile alignments identify syntenic homologs outside Saccharomyces for 19 genes identified as SRGs in BLAST searches. A) Negative log e-values of hh-suite profile-profile alignments between MSAs of SRGs and MSAs of groups of homologous genes found in genomes outside Saccharomyces. Three groups are considered: Genus-conserved SRGs vs. homology groups constructed from genes that share synteny with the SRG, Genus-unconserved SRGs vs. homology groups constructed from genes that share synteny with the SRG, and, as negative controls, genus-conserved SRGs vs. homology groups constructed from genes that share synteny with some other genus-conserved SRG selected at random. The dashed line gives the 5% FDR threshold. B) Distribution of species with potential candidate homologs (an ORF > 300 bp) of SRGs in syntenic regions in post-WGD genomes outside Saccharomyces. Genus-conserved SRGs are much more likely than negative controls to have candidate orthologs among at least 50 species (odds ratio 6.64, p=.001, Fisher’s exact test), while genus-unconserved SRGs are not (odds ratio 1.15, p=1), C) For each SRG with inferred syntenic homologs outside Saccharomyces, the proportion of these syntenic homologous that are located on the expected strand given the relative strand orientation of the SRG and anchor gene in S. cerevisiae. D) The ORF length and percentile of Saccharomyces amino acid divergence is given for inferred vertically descended SRGs and 100 non-SRG genes selected at random. The amino acid divergence of a gene is average pairwise alignment distance between the encoded proteins of each pair of orthologs of that gene within the Saccharomyces genus. The median length of single-exon non-SRG annotated genes (1296 nt) is indicated by a dashed line.

To determine whether the lack of identified homologs for genus-unconserved SRGs was due to lack of statistical power, we evaluated whether, among SRG ORFs larger than 300 bp, there was an excess of ORFs >300 bp in extrageneric syntenic regions that could be potential homologs. Many genus-conserved SRGs, but not genus-unconserved SRGs, show potential syntenic homologs in a large majority of extrageneric syntenic regions indicating that the difference in homolog detection using hh-suite is not simply due to statistical power (Figure 3B).

Of the 19 SRGs with syntenic matches, 14 also have identified homologs in Yeast Genome Order Browser (YGOB)^27^ (Supplementary Figure 3), indicating high concordance between our automated, statistically controlled approach and the manual curation performed by YGOB. No genus-unconserved SRGs have identified homologs on YGOB. In our approach, the use of anchor genes to identify candidate syntenic homologs provides an additional opportunity for validation of the vertical descent relationships inferred by profile-profile alignments. If the anchor and SRG are on the same strand in *Saccharomyces*, we also expect them to usually be on the same strand outside *Saccharomyces*, and vice versa. For 18 of the 19 SRGs with syntenic matches, the strand agrees with the prediction in most non-*Saccharomyces* genomes that contain the inferred homologous sequence (Figure 3C; inferred YGL015C homologs are opposite their expected orientation). These data provide strong orthogonal support to our inferences of vertical descent between SRGs and their divergent syntenic homologs.

Of the remaining 20 genus-conserved SRGs for which we did not identify syntenic homologs, only one has identified homologs in YGOB: YPL200W, a paralog of YGL075C derived from the WGD.^27^ This gene appears to have been lost independently several times post-WGD and was missed by our approach because we only looked for syntenic homologs in post-WGD species.

To better understand why BLAST failed to find homologs that did exist, we examined the properties of the 20 SRGs inferred to have homologs outside *Saccharomyces* with preserved synteny (19 identified by our automated pipeline plus YPL200W). The proteins encoded by these genes tend to have a high level of divergence within *Saccharomyces*, indicative of rapid evolutionary rates (Figure 3D). These vertically-descended SRGs are also relatively short, with all 20 below the median length of 1299 nt among all genus-conserved ORFs in *S. cerevisiae* (Figure 3D). Both short length and rapid divergence pose challenges for homology detection.^12^

Given our observations, we suspected that some SRGs derived by vertical descent may be evolving too quickly to be detected even by our automated profile-profile alignment approach. Through manual evaluation of syntenic regions and other evidence we identified six additional SRGs that showed strong evidence of vertical descent (Appendix 1, Supplementary Figure 4-6). Overall, our analyses reveal that 26 of 39 genus-conserved SRGs likely entered the *Saccharomyces* genome by vertical descent, with strong matches to likely homologs outside *Saccharomyces,* while we find no evidence that any genus-unconserved SRGs derive from vertical descent.

### SRGs are often associated with transposable elements

Transposable elements (TEs) are a potential source of TRGs due to their ability to transfer horizontally between distantly related genomes.^28,29^ To determine whether any SRGs are derived from transposition events, we assessed whether SRGs were enriched in TE hotspots, as indicated by being located within 1kb of an annotated TE. Expectedly, we observed no significant enrichment near TEs relative to annotated genes overall among groups of SRGs for which we had already found support for non-TE origins: the 169 SRGs for which parsimony supports *de novo* birth (p=0.63, binomial test) and the 26 SRGs we inferred to be derived by vertical descent (Figure 4A; p=0.85, binomial test). Among the remaining unexplained SRGs, 18 of 102 SRGs lacking genus-level conservation are within 1 kb of a TE, a 3.4-fold enrichment relative to annotated genes overall (p=4.3 × 10^−6^, binomial test), while 5 of 12 genus-conserved SRGs (42%) are within 1 kb of a TE, an 8.1-fold enrichment (p=2.1 × 10^−4^, binomial test, Figure 4A). We can estimate the approximate proportion of SRGs with transposable element associated origins from the difference between the frequency of TE-adjacency among SRGs and the frequency among genes overall. This analysis suggests that approximately 13 of 271 genus-unconserved SRGs (5%), and 5 of 39 genus-conserved SRGs (13%), are of TE origin. This likely represents an underestimate as TE-derived genes can translocate away from the hotspot after transposition.

**Figure 4:**
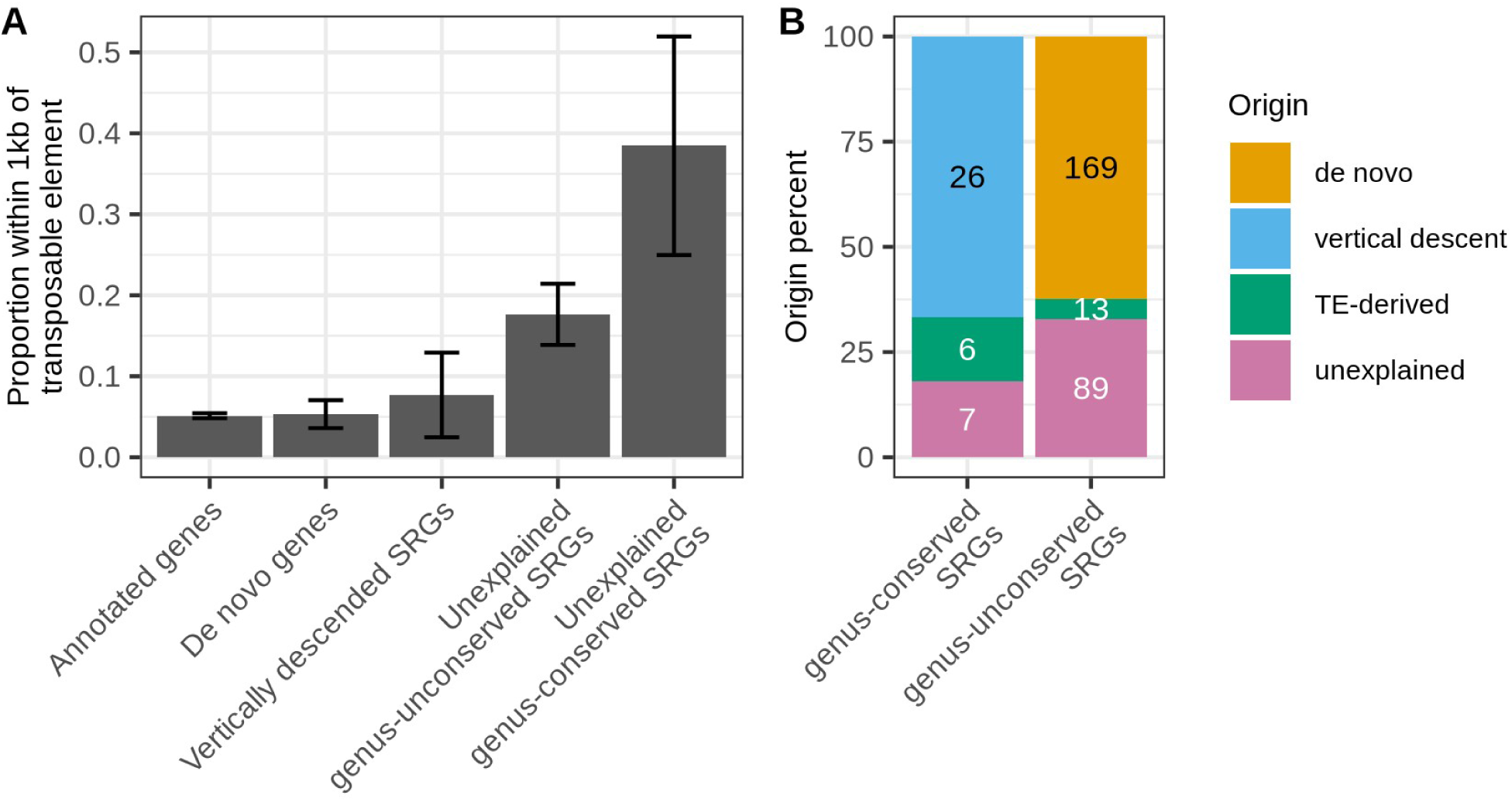
Transposable element associated mechanisms explain the origins of some of SRGs. A) Among several groups of genes, the proportion of genes in the group within 1 kb of an annotated transposable element feature on SGD. Annotated genes consist of all single-exon genes annotated on SGD (N=5581). De novo genes are genes with de novo origin supported by parsimony (N=169). Vertically descended genes (N=25) are genes with identified homologs outside Saccharomyces that appear to be related to the Saccharomyces gene by vertical descent. Unexplained SRGs (N=102 genus-unconserved, N=13 genus-conserved,) are SRGs which were not classified as de novo or of vertical descent. B) Inferred origins for genus-conserved and genus-unconserved SRGs. Counts for de novo and vertical descent are in black to indicate that each inferred count in these categories comes from assigning this origin to a specific individual gene. Counts for TE-derived origins, in white, are estimated from enrichment of SRGs as a group in TE hotspots.

We next searched for evidence of homologs outside Saccharomyces that could provide additional support for a TE-related mechanism of origin. We found that the gene YIL092W, a genus-conserved SRG located outside TE hotspots, appears to be derived from a TE Gag gene, while two other genus-conserved SRGs located in TE hotspots have a phylogenetic distribution of homologs outside the genus consistent with TE origin (Appendix 2, Supplementary Figure 7). Adding YIL092W to the five genus-conserved SRGs in TE hotspots, we thus estimate that at least six genus-conserved SRGs entered the *Saccharomyces* genome from a TE-associated mechanism, likely horizontal transfer (Figure 4B, Supplementary Table 1).

### Resolving the origins of SRGs provides clarity on the evolutionary role of *de novo* gene birth

Of 39 genus-conserved SRGs, we found that 26 are likely derived from vertical descent and an additional six are strong candidates by origination via TE-mediated horizontal transfer (Figure 4B). This would leave only seven for potential origins in *de novo* gene birth. This result contrasts with most previous studies of *de novo* gene birth in *S. cerevisiae*, which have reported many more *de novo* gene candidates that are conserved across *Saccharomyces.*^7,17,18,30,31^ The most recent such study, Montañés et al. 2023^18^, identified 300 S. *cerevisiae* genes as “putative *de novo* genes” that had no homologs outside *Saccharomyces*, of which 94 are classified as genus-conserved in our analysis. This amounts to over thirteen times more putative *de novo* genes conserved across *Saccharomyces* than the seven genus-conserved *de novo* candidates remaining in our analysis.

To better understand the source of this discrepancy, we classified each of the 94 genus-conserved SRGs from Montañés et al. 2023^18^ by the manner in which we accounted for their origin in our own analysis (Figure 5A). The largest source of discrepancy is that we identified homologs outside *Saccharomyces* for 62 of the 94 (66%) genes in our initial BLAST search. This is presumably explained by our use of a much larger set of comparison genomes than Montañés et al. 2023.^18^ For an additional 20 (21%) genes, initial homology searches failed to identify homologs outside *Saccharomyces* in either analysis, but we found support for vertical descent using profile-profile alignments. Thus, use of a larger collection of comparison genomes and sensitive profile-profile alignment methods enables rejecting recent *de novo* origins for most genus-conserved genes identified as putative *de novo* genes in prior work.

**Figure 5.**
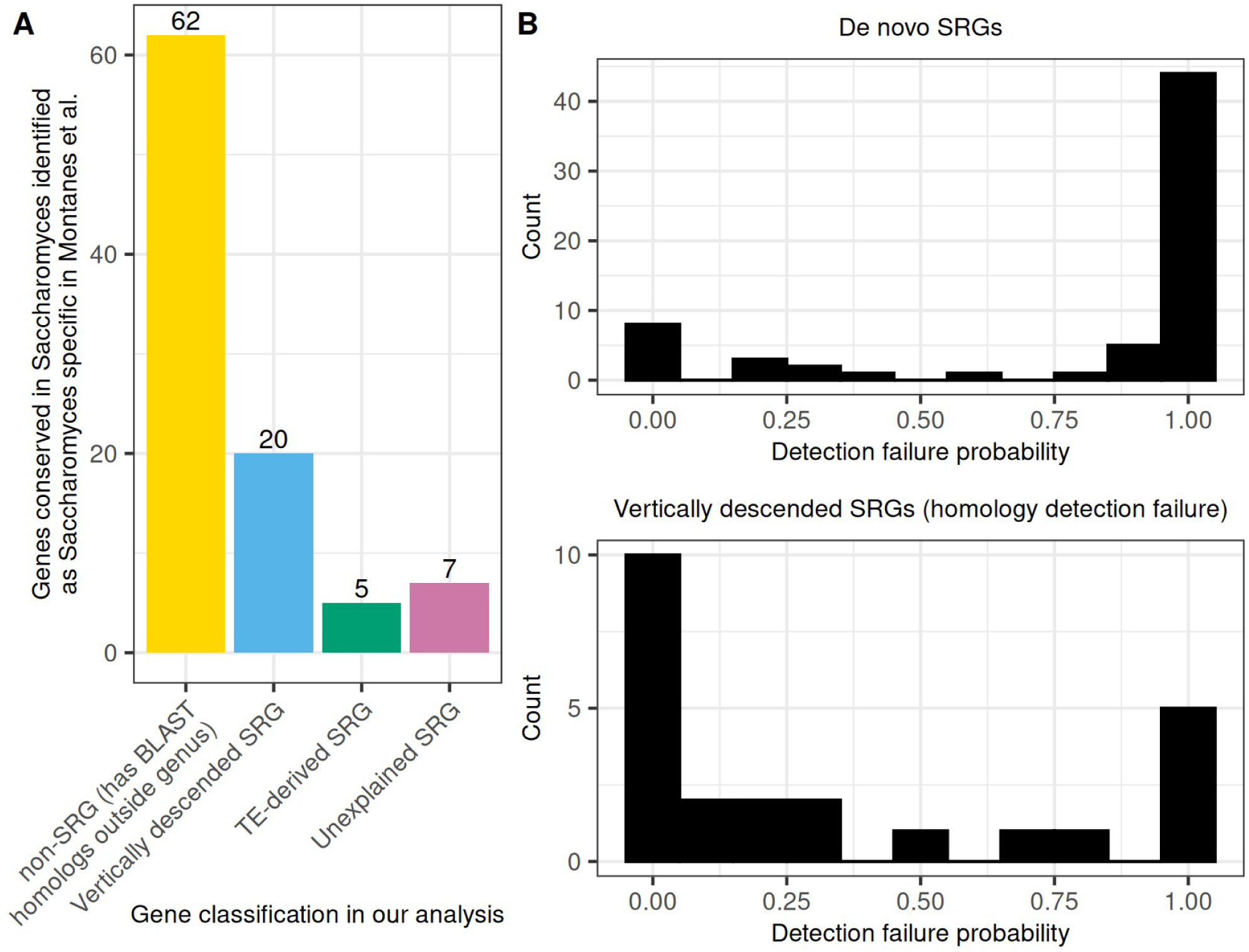
Resolving the origin of SRGs provides clarity on the role of de novo gene birth in evolution. A) Among Saccharomyces-conserved genes inferred to have originated “putatively de novo” since Saccharomyces-Nakaseomyces divergence in Montañés et al. 2023^18^ (i.e, putative Saccharomyces-specific genes), we indicate the inferred classification from our own analysis conducted in this study. B) Detection failure probabilities indicated in Weisman et al. 2020^12^ for SRGs identified as either of de novo origin (top) or vertical descent (bottom) in this study.

Given our ability to assign origins to previously ambiguous cases, we next assessed how our results affect the controversy over whether the prevalence of TRGs is better explained by *de novo* gene birth or homology detection failure. Vakirlis et al. 2020^15^ used a statistical model based on the presence of candidate homologs for SRGs in syntenic regions to estimate that at most 45% of SRG*s* are explained by sequence divergence of homologs beyond recognition by BLAST, as opposed to *de novo* gene birth or horizontal transfer. Our result is consistent with the Vakirlis et al. 2020 prediction as we find that only 26 of 310 SRGs (8.4%) show evidence of having homologs missed by BLAST due to sequence divergence (excluding horizontal transfers). Our results thus broadly support the proposition that TRG prevalence is explained primarily by frequent *de novo* gene birth.

Another influential paper, Weisman et al. 2020^12^, developed the abSENSE model to estimate the probability that BLAST would fail to detect a SRG in a comparison genome. These authors found that a majority of SRGs are evolving sufficiently rapidly that the model predicts a failure to detect any existing homologs outside the genus, and so lack of such detected homologs provided no evidence the gene derives from *de novo* gene birth. To understand how our results relate to this model, we obtained the homology detection failure probabilities calculated by abSENSE for all SRGs we analyzed. Unexpectedly, among genes we are able to assign to either *de novo* origin or homology detection failure, we find that *de novo* SRGs tend to show much higher susceptibility to homology detection failure than the SRGs we demonstrate actually are explained by homology detection failure (Figure 5B; p=9 × 10^−6^ , permutation test for difference in mean). This apparent paradox is explained however. The genes most statistically susceptible to homology detection failure are short and fast evolving, as small size and rapid divergence pose challenges for homology detection algorithms. True *de novo* genes also tend to be among the shortest and fastest evolving genes in the genome (Supplementary Figure 8). As a result, abSENSE assigns them high probability for homology detection failure. Weisman et al. 2020 are correct that it is logically invalid to infer likely *de novo* origin merely from the observation that a small and rapidly evolving gene lacks homologs outside its close relatives—its lack of detected homologs is expected even if it derives from vertical descent—but a high probability for homology detection failure also does not invalidate *de novo* origin since the shortest and fastest evolving genes tend to be *de novo* in any case.

In sum, we show that most genus-conserved SRGs proposed as candidate *de novo* genes in prior work are likely explained by vertical descent or horizontal transfer rather than *de novo* origin. Nevertheless, *de novo* origin remains the primary explanation for SRGs overall because it is the dominant origin for the largest class of SRGs, those restricted below the genus level.

## Discussion

Whether the prevalence of TRGs across all life and all taxonomic ranks reflects widespread *de novo* gene birth or technical limitations in homology detection algorithms is crucial for understanding the broad evolutionary significance of *de novo* gene birth. Here, we find strong support for *de novo* gene birth as the primary explanation for genes restricted to the *Saccharomyces* genus (SRGs). This result is driven by SRGs that lack genus-level conservation, which make up a large majority of all SRGs. Most genus-unconserved SRGs can be shown to be of *de novo* origin through multiple sequence alignments of closely related species, whereby an ancestral noncoding ancestor can be inferred to have existed prior to emergence of the gene.^31^ Moreover, even genus-unconserved SRGs for which *de novo* origin cannot be directly inferred from an alignment show signs that they are not explained by homology detection failure: sensitive profile-profile alignments fail to improve on BLAST in identifying homologs for these genes in syntenic regions outside the genus, and these genes also show no enrichment above null expectations for the presence of candidate syntenic homologs that could be too diverged for recognition by profile-profile alignment. *De novo* gene birth thus appears to be by far the dominant mechanism explaining *S. cerevisiae* genes restricted below the genus level, with some contribution from TE-mediated horizontal transfer.

Yet, for SRGs that are conserved at the genus level, concerns about homology detection failure appear well-founded.^12^ We find that most genus-conserved SRGs have syntenic homologs identifiable through sensitive profile-profile alignments, and others are strongly enriched in transposable element hotspots^32^, suggesting origins by a TE related mechanism such as horizontal transfer.^28^ Even among seven genus-conserved SRGs that remain unexplained, we have no reason to expect these are primarily of *de novo* origin. Our homology detection approach depends on both the maintenance of synteny and the preservation of some degree of sequence similarity, and so will miss homologs of genes that have lost synteny or diverged even more quickly than the genes whose homologs we are able to recover. Thus, *de novo* birth appears at most a minor contributor to SRGs that are conserved across the genus.

The overall implication of our findings for the quest to determine the mechanisms of gene origination, then, is that species-specific SRGs are mostly unaffected by homology detection failure while SRGs conserved across the genus are mostly explained by it. Thus, the difference in magnitude between the number of *de novo* species-specific genes and older *de novo* genes appears starker than in prior analyses on either side of the controversy. Moreover, our study undercounts the number of very young *de novo* genes because we consider only ORFs annotated in the leading genome database SGD. Ribosome profiling studies show that thousands of short unannotated yeast sequences that are not conserved throughout the *Saccharomyces* genus are also translated^4,33^ and can encode microproteins that provide benefits to the organism.^4,34,35^ Based on our findings, these short unannotated translated sequences are unlikely to be affected by homology detection failure and are expected to overwhelmingly be of *de novo* origin. Together, these observations imply strong filters that prevent almost all *de novo* genes from persisting over deep evolutionary time.

It is of interest to determine whether the patterns we report among yeast TRGs hold more broadly across life. For example, some of the best characterized TRGs, such as *Goddard* and *Saturn* in *Drosophila*, are often referred to as “putative *de novo* genes” as they have not yet been either proven or disproven to be of *de novo* origin.^36^ The methods we developed here could be adapted into a general strategy for resolving TRGs as either of *de novo* origin or explained by homology detection failure. As yeast genera tend to be much more genetically diverse than the genera of plants or animals^22^, we expect that even if the general evolutionary dynamic is similar across life, patterns in particular taxonomic ranks like “genus” will differ across taxa. Genes with strong evidence of *de novo* emergence, almost all very young, are being characterized at a rapid pace across taxa, revealing diverse functions.^37–39^ Whether *de novo* genes ever persist over deep evolutionary time, or are restricted to evolutionarily transient roles, remains a key open question.

Overall, our results enable reconciliation of prior conflicting models and provide clarity on what information TRGs can provide about *de novo* genes. *De novo* gene birth appears very common in *Saccharomyces*. As a result, there is an abundance of *de novo* genes genuinely restricted to low taxonomic ranks, and these make up an absolute majority of SRGs. On the other hand, *de novo* gene persistence appears quite rare. Thus, as we climb taxonomic ranks, the signal of genuine *de novo* genes declines while the noise of homology detection failure persists, and our ability to do evolutionary inference correspondingly weakens (Figure 6).

**Figure 6.**
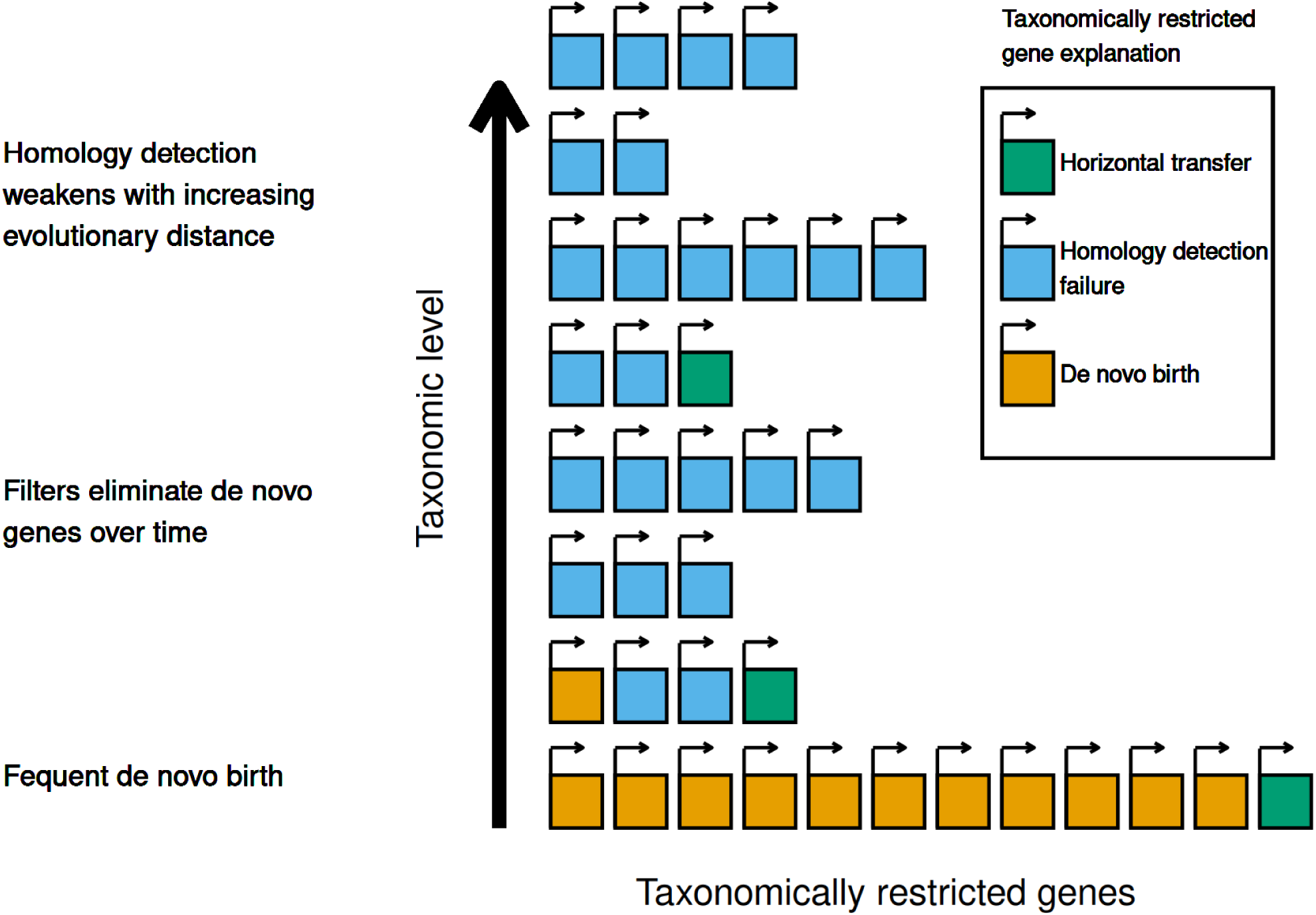
A model of mechanisms underlying the prevalence of taxonomically restricted genes. Conceptual illustration of a model proposed to explain the prevalence of TRGs across different taxonomic levels. The lowest taxonomic level is dominated by frequent de novo gene birth, but strong filters prevent almost all de novo genes from persisting through deep evolutionary time. With low rates of de novo gene persistence, few TRGs at higher levels are survivors of past de novo gene events. Instead, patterns of taxonomic restriction at these ranks are primarily explained by homology detection failure, as homology detection becomes increasingly difficult with evolutionary distance. Horizontal transfer contributes TRGs at all levels but at low rates.

## Methods

### Identification of ***Saccharomyces***-restricted genes

We attempted to identify taxonomically-restricted genes among *S. cerevisiae* annotated genes. All information on annotated genes was taken from the Saccharomyces Genome Database^40^ dated September 17, 2025. We considered a gene to be annotated if it was classified as “verified” or “uncharacterized” in Saccharomyces Genome Database. A FASTA file consisting of the protein sequences of each such gene was constructed. As a negative control, we also randomly scrambled each sequence in the *S. cerevisiae* proteome and appended these sequences to the file. We used BLASTP^41^ and TBLASTN to query these protein sequences and controls against the protein sequences (for BLASTP) or whole genomes (for TBLASTN) for each of the 1,146 genomes assembled in Opulente et al. 2024^21^ from species in the subphylum *Saccharomycotina*. Default BLAST settings were used except for setting an e-value threshold of .01.

We took steps to guard against the possibility of contamination in the Opulente et al. 2024 dataset creating false inferences of homology. For any BLASTP or TBLASTN hit, we extracted the nucleotide sequence in the *Saccharomycotina* genome corresponding to the matched sequence and queried it against the *S. cerevisiae* nucleotide sequence using BLASTN. Matches with nucleotide identifty > 90% and e-values < 10^−4^ against any *Saccharomyces* genus genome were considered potential contamination and not used to infer homology.

Genes were considered to have homologs outside *Saccharomyces* (and thus not be *Saccharomyces*-restricted) if they met one of three criteria: 1) BLASTP or TBLASTN match with e-value < 10^−4^ with at least three species outside the *Saccharomyces* genus 2) BLASTP or TBLASTN match with e-value < 10^−6^ with at least one species outide the *Saccharomyces* genus 3) e-value < 10^−6^ against an *S. cerevisiae* gene that met one of the first two criteria. These thresholds were set based on the minimal number of negative controls passing them, as shown in Supplementary Figure 1B. All *S. cerevisiae* genes lacking homologs outside *Saccharomyces* by these criteria were considered *Saccharomyces*-restricted genes (SRGs).

### Distinguishing genus-level conservation and *de novo* inference

Reading frame conservation (RFC) values for each *S. cerevisiae* gene were taken from Wacholder et al. 2023.^4^ These data provide the RFC between the *S. cerevisiae* ORF and its inferred ortholog in each other *Saccharomyces* species, or an indication that no ortholog was identified in that species. Following Wacholder et al. 2023, genes were considered conserved in the genus if they had mean RFC > 0.8 among identified orthologs in the genus.

Inference of *de novo* origin was based on patterns on ORF presence/absence among the species of *Saccharomyces*. ORF presence was inferred if the pairwise RFC between the *S. cerevisiae* gene and its ortholog in the species exceeded 0.8. *De novo* origin is supported by parsimony if the ORF is absent in *S. uvarum*, *S. eubayanus*, and *S. arboricola*. ORF presence or absence was only inferred if the orthologous region with identifiable nucleotide-level homology was identified in Wacholder et al. 2023.^4^

### Detection of syntenic homologs by profile-profile alignment

Detection of syntenic homologs involved several steps, illustrated in Supplementary Figure 2. First, for each SRG in the *S. cerevisiae* genome, the three closest annotated genes in each direction were selected as anchors. Homologs of these anchor genes among the Opulente et al. 2024^21^ post-WGD genomes (excluding *Saccharomyces* genomes) were identified using BLASTP against the protein list from that paper, with a 10^−4^ e-value threshold. For the anchor homologs in each comparison genome, all ATG-to-stop ORFs within 10kb were identified as candidate syntenic homologs of the SRG.

The candidate syntenic homologs were then clustered into homology groups according to the following process: An all-vs-all BLASTP search was conducted among all candidates. The candidate with the most BLASTP hits (at a 10^−4^ e-value threshold) among other candidates was selected, and that candidate together with all hits were taken to form a homology group. The number of hits of each remaining candidate was then recalculated, excluding the candidates already assigned to a homology group. Then the process was repeated until either all candidates were assigned to a homology group or the remaining candidates had no BLASTP matches among themselves.

Within each homology group, the candidates were aligned into an MSA using the Muscle5 sequence aligner.^42^ The hhalign program from hh-suite^26^ was used to construct a profile from the MSA. An MSA and profile was also constructed for the SRG within *Saccharomyces*, using the *S. cerevisiae* sequence together with its orthologs identified in Wacholder et al. 2023.^4^ Profile-profile alignment between the *Saccharomyces* MSA and each homology group MSA was then conducted using the hhalign program from hh-suite. This provides an e-value, which was used as an indicator of the strength of evidence for homology in downstream analyses.

In addition to the tests between MSAs of each SRG and that of its candidate homolog homology groups, as a negative control we also tested each SRG against the candidate homolog homology groups of a different, randomly selected SRG. In this test, genus-conserved SRGs were matched against the candidate homologs of other genus-conserved SRGs, and genus-unconserved SRGs were matched against those of other genus-unconserved SRGs. We have no reason to expect genuine homology in these comparisons, so these tests provide an indication of the null distribution of e-values when no homolog is present.

### Detection of enrichment for candidate syntenic homologs

In the approach outlined above for detecting specific homologs of SRGs, detection sensitivity depends on protein length and divergence from the SRG. We also developed a modified approach to identify plausible candidate homologs regardless of whether we could confirm homology using profile-profile alignment.

In this approach, anchors were selected and anchor homologs identified in the same manner as described above. However, we required anchors within 10 kb of each candidate ORF in both directions, as a single-anchor approach would generate too many potential homologs. Any start-to-stop ORF >300 bp was then considered a candidate homolog.

### Properties of identified syntenic homologs

To assess whether SRGs had reported homologs in Yeast Genome Order browser, we downloaded the Pillars.tab table from the Yeast Genome Order Browser website Version 7.^27^ We then checked whether each SRG had any gene sharing the same pillar (indicating inferred homology) in genomes outside *Saccharomyces*.

To calculate percentile of amino acid divergence for each *S. cerevisiae* gene, we first constructed a multiple sequence alignment of the protein sequence encoded by the gene together with that of its orthologs identified in Wacholder et al. 2023^4^ using the Muscle5 sequence aligner.^42^ We then calculated the average pairwise distance within the MSA using the dist.alignment function from the seqinr R package^43^ and converted these values to percentiles across annotated genes.

### Identifying homologs of transposable elements

To find potential homologs of transposable elements, we first extracted the protein sequences of all transposable element genes annotated on SGD. We then identified homologs of these transposable elements by using TBLASTN and BLASTP of each sequence against the yeast genomes and proteomes collected in Opulente et al. 2024, adding each match with e-value < 10^−4^ as a potential homolog.^21^ We then constructed homology groups among all TEs and their homologs, produced an MSA of all protein sequences within each homology group, and created a hh-suite profile from the MSA as described above. We then performed profile-profile alignments of each SRG against each TE homology group profile and extracted the e-values associated with these alignments. Since some TE homology groups were very large, we randomly selected 200 sequences within each group to generate the MSA and perform subsequent analyses if the group had more than 200 total sequences.

### Identifying genes located in transposable element insertion hotspots

We identified TE insertion hotspots as regions of the S. cerevisiae genome near transposable element annotations. We extracted the coordinates of all genomic elements annotated as “long terminal repeat”, “LTR retrotransposon” or “transposable element gene” from SGD. For each protein-coding gene, the distance of its CDS to the nearest TE annotation was calculated.

### Comparison to prior work on SRGs

Data from Montañés et al. 2023^18^ was extracted from Supplementary Table 2 of that publication. A gene was considered to have arisen “putatively de novo” after *Saccharomyces* divergence (and thus be *Saccharomyces*-specific) if it was assigned to age nodes N0 through N4. Genes were considered to be genus-conserved according to the definition used in our analyses; i.e., if average reading frame conservation between the *S. cerevisiae* ORF and its orthologs exceeded 0.8. Homology detection failure probabilities for *S. cerevisiae* genes calculated by Weisman et al. 2020^12^ were taken from the GitHub page for abSENSE.

## Appendix 1: Identification of vertically descended SRGs by investigation of individual cases

### YBR063C is a rapidly evolving gene with many likely syntenic homologs among related genera

We assessed whether there were any good candidates for SRGs derived by vertical descent that were evolving too quickly to be detected even by our automated profile-profile alignment approach. We manually examined syntenic regions for each SRG in genomes outside *Saccharomyces*. We found two strong candidates, YBR063C and YJL118W. YBR063C is large for an SRG at 1215 nt and extremely fast evolving within the *Saccharomyces* genus, above the 99^th^ percentile of intra-*Saccharomyces* amino acid divergence of *S. cerevisiae* genes. On YGOB^27^, representative species from six different genera have genes in the syntenic region of YBR063C between YBR061C and YBR065C that are not assigned as homologs to genes in any other genera. This pattern is expected if YBR063C evolves so quickly that its orthologs cannot be recognized between different genera. To test this hypothesis more formally, we identified all ORFs longer than 300 nt in the syntenic block between YBR061C homologs and YBR065C homologs in all post-WGD species; this block is well-preserved in post-WGD genera. We identified 35 post-WGD genomes with at least one ORF longer than 300 nt in this syntenic block; most had only a single such ORF (Supplemental Figure 4A). We then clustered these ORFs into homology groups independently for each genus and performed profile-profile alignment of each homology group against the *Saccharomyces* YBR063C MSA using hh-suite. The YBR063C MSA matched two *Kazachstania* homology groups with e-value <.01 (e-values .0039 and .0066); these groups together represent the sole ORF >300 nt in the syntenic blocks of 11 *Kazachstania* species and all are in the expected orientation for a YBR063C ortholog (Supplemental Figure 4A-B). The YBR063C MSA did not match to homology groups from other genera directly; however, groups consisting of genes from each genera showed evidence of homology with some other groups (e-value <.01), ultimately connecting the genes across seven genera as likely YBR063C orthologs related by vertical descent (Supplemental Figure 4B). Altogether, these results indicate that YBR063C is a consistently fast evolving gene, with its orthologs rapidly diverging past the threshold of recognition by BLAST.

### YJL118W appears to be a highly diverged ohnolog of YKR046C

The second candidate for origin by vertical descent, YJL118W, is 660 nt and slow evolving for an SRG, at the 29^th^ percentile of amino acid divergence in *Saccharomyces*. In *S. cerevisiae*, YJL118W is located between YJL117W and YJL121W. According to YGOB^27^, synteny between YJL117W and YJL121W predates the WGD event and it is maintained in post-WGD genera with the exception of *Kazachstania*. Also according to YGOB, three genes were located in between YJL117W and YJL121W in the ancestral locus, homologous to the *S. cerevisiae genes* YKR045C, YKR046C, and YKR048C. The locus with all five genes is maintained in some post-WDG lineages, such as the *Zygosaccharomyces* genus, but in *Saccharomyces* YJL118W and YJ121W survived on one copy of the duplicated locus (now on chromosome X) while YKR045C, YKR046C, and YKR048C survived on the other copy (now on chromosome XI) (Supplemental Figure 4C).

These observations suggests the hypothesis that YJL118W could be a highly diverged ohnolog (i.e., a paralog resulting from a WGD^44^) of YKR045C, YKR046C, or YKR048C. The most plausible candidate is YKR046C, because homologs of this gene identifiable by genome-wide BLASTP are still present between YJL117W and YJL121W and between YKR045C and YKR048C in *Tetrapisispora* and *Vanderwaltozyma* species (Supplemental Figure 4C, Supplemental Figure 5). This indicates that both copies of YKR046C originating from the WGD still existed in the common ancestor of *Saccharomyces*, *Kazachstania*, *Tetrapisispora* and *Vanderwaltozyma*. However, the *Saccharomyces* YJL118W MSA has no significant similarity by profile-profile alignment to the *Saccharomyces* MSA of YKR046C (e-value >.05).

In *Kazachstania*, synteny between YJL117W and YJL121W has been lost; instead YJL121W is near a homolog to YBR103W (Supplementary Figure 6). For most Kazachstania species, between one and two ORFs longer than 300 nt are located between YJL121W and YBR103W homologs, with neither identifiably homologous to any *S. cerevisiae* gene by genome-wide BLASTP at a 10^−4^ e-value threshold. We clustered these ORFs into homology groups. The largest *Kazachstania* homology group (Group 1 in Supplementary Figure 4C) has a match to both YJL118W (e-value = .0021) and YKR046C (e-value = 10^−10^) by profile-profile alignment. Taking this match into consideration together with a phylogenetic history of the relevant loci after the WGD suggests a plausible evolutionary account (Supplemental Figure 4C). In the lineage leading to the common ancestor of *Kazachstania* and *Saccharomyces*, one copy of YKR046C diverged such that it is not identifiable as a YKR046C homolog by BLASTP, but can be identified as such by more sensitive profile-profile alignment. In *Saccharomyces*, this copy (YJL118W) diverged even further than in *Kazachstania* such that it is not identifiable as a homolog to *Saccharomyces* YKR046C even by profile-profile alignment, but still matches by profile-profile alignment to its *Kazachstania* ortholog, which it shares the most evolutionary history with of any extant homolog outside *Saccharomyces*. Thus, YJL118W appears to be an ohnolog of YKR046C. This case illustrates the challenges of confidently establishing vertical descent for SRGs that have greatly diverged from their close orthologs even when synteny is maintained.

### Profile-profile alignment facilitates detection of functionally related metallothionein orthologs of CUP1-1 and CUP1-2

Just as synteny can increase the power of homology detection by limiting the number of possible candidates, functional similarity between potential homologs can do the same. Unfortunately, most SRGs are poorly characterized. A major exception are the genes CUP1-1 and CUP1-2, which are identical and adjacent in the S288C reference genome. These genes encode 61 aa copper metallothioneins, proteins that bind copper in order to prevent toxicity.^45^ Copper metallothionines are cysteine-rich proteins that have been extensively studied in many species, including *Nakaseomyces glabrata*, a human pathogen in a closely related genus to Saccharomyces.^46^ Though characterized metallothionines are highly diverse at the sequence level and are not obviously recognizable as homologs, some researchers have proposed that multiple yeast metallothionines form a gene family including both CUP1 and the *N. glabrata* metallothionine MT1 on the basis of apparent conservation of several cysteine residues.^47^ *S. cerevisiae* CUP1 and *N. glabrata* MT1 are also regulated by homologous transcription factors and in a similar manner.^48^ Pairwise BLASTP between MT1 and CUP1 gives an e-value of 0.001, which is strong if treated as a pairwise test but would be insufficient to infer homology in a genome vs. genome comparison.

To further assess whether *S. cerevisiae* CUP1 is homologous to other yeast metallothionines, we performed profile-profile alignment of CUP1 against metallothionines in each other post-WGD genus. On YGOB, CUP1 has no syntenic homologs with species outside *Saccharomyces*. However, *N. glabrata* MT1 has syntenic homologs in several other genera. As MT1 is nearby a homolog of well-conserved *S. cerevisiae* gene YPR079W, we used YPR079W as an anchor gene in a similar manner to our previous analyses: identifying all ORFs within 10 kb of each YPR079W homolog in each genome, clustering these ORFs into homology groups within each genus, and then performing profile-profile alignment using hh-suite for each homology group against CUP1. CUP1 showed strong pairwise profile-profile e-values with homology groups sharing synteny with MT1 in *Nakaseomyces* (4 × 10^−5^), *Kazachstania* (1.9 × 10^−4^), *Vanderwaltozyma* (1.2 × 10^−7^) and *Zygosaccharomyces* (9.5 × 10^−6^). Notably, the strong alignment with the *Vanderwaltozyma* MT1 homolog covers almost the entirety of CUP1 and five cysteine residues appear fully conserved between the genera (Supplemental Figure 4D). These results indicate that the BLASTP e-value is not spurious and *Saccharomyces* CUP1 has deep sequence similarity to metallothionines in related genera.

We cannot completely rule out the possibility that *Saccharomyces* CUP1 is derived from *de novo* birth and the sequence similarity is due to convergent evolution. However, the plausible evolutionary scenario of vertical transfer is simple: the CUP1/MT1 gene was present at the same locus for tens of millions of years and then experienced a translocation in the *Saccharomyces* lineage, perhaps together with its promoter, that disrupted synteny with its homologs in other genera. In a *de novo* birth scenario, the *Saccharomyces* lineage must have lost its ancestral metallothionine only to independently evolve a new gene performing the same function, with the same regulation, by a sufficiently similar mechanism to obtain highly significant sequence similarity to metallothionines in related species. Moreover, we have already seen that the great majority of genes conserved within *Saccharomyces* entered the *Saccharomyces* genome through vertical descent and not *de novo* birth, so even if each scenario were otherwise equally plausible vertical descent should be favored. We conclude that it is of very high likelihood that *Saccharomyces* CUP1 is derived from vertical descent and is homologous with *N. glabrata* MT1.

### SRG YLR125W is a paralog of non-SRG YMR030W

Another potential source of TRGs is gene duplication followed by rapid divergence that causes homology detection failure. Just as profile-profile alignment can detect syntenic orthologs more sensitively than BLAST, it can also facilitate identifying highly diverged paralogs. To identify potential SRGs that underwent duplication followed by divergence, we searched for paralogs of each SRG within the *S. cerevisiae* genome. We conducted profile-profile alignments between the MSAs of each SRG and the MSAs of each annotated non-SRG gene in *S. cerevisiae*, with each MSA consisting of the gene and its *Saccharomyces* orthologs.

One genus-conserved SRG, YLR125W, has a single strong profile-profile alignment match to YMR030W (e-value 2.9 × 10^−8^), a non-SRG with 11 BLASTP matches to genomes outside *Saccharomyces* (e-value < 10^−4^) mostly in *Kazachstania* and *Naumovozyma*. YMR030W (standard name RSF1) appears to be a transcription factor necessary for respiratory growth^49^ while YLR125W is uncharacterized. The protein encoded by YMR030W has only three physical interactions listed on SGD, one of which is to YLR125W, identified in a two-hybrid screen by Ito et al. 2001.^50^ The physical interaction between these paralogs suggest the possibility that the YMR030W/YLR125W ancestor was a homodimer and a gene duplication led to heterodimerization. Overall, these observations are consistent with a duplication-divergence scenario and indicate that YLR125W did not emerge *de novo* in *Saccharomyces*.

### Homologs of YEL073C outside ***Saccharomyces*** identified by searching using ***S. uvarum*** sequence

The genus-conserved SRG YEL073C encodes a protein of 107 amino acids. We observed that the YEL073C homologs in *S. uvarum* and *S. eubayanus* were extended on the 3’ end relative to *S. cereivisae*, predicted to encode proteins of 145 and 141 amino acids, respectively. Given the challenges in homology detections for small proteins, we hoped that using the larger protein sequences in these species as queries might enable detection of homologs outside *Saccharomyces*. We therefore used BLASTP to query the *S. uvarum* sequence against the yeast genomes assembled by Opulente et al. 2024.^21^ We found strong matches (e-value < 1 × 10^−6^) among nine species in the *Zygotorulaspora*, *Torulaspora*, *Lachancea*, *Kazachstania*, and *Suhomyces* genera, including six very strong matches (e-value < 1 × 10^−9^).

## Appendix 2: Evidence for transposable element origins for individual SRGs YIL092W is derived from a Gag gene

To identify genes potentially derived from core transposable element genes, we constructed homology groups for all annotated TE genes on SGD and their homologs in genomes collected by Opulente et al. 2024.^17^ We then performed profile-profile alignments of these TE homology groups against each SRG. Additionally, we submitted each SRG to the Censor tool^51^, which searches for homology within the Repbase database of repetitive elements. A single 1902 nt ORF, YIL092W, matched a homology group containing the *S. cerevisiae* TE genes YGR109W-A and YIL082W together with 272 homologous genes in other species (e-value = 3.28 × 10^−9^). YGR109W-A and YIL082W are both annotated as Gag genes from the Ty3-*gypsy* group of LTR retrotransposons. The same YIL092W ORF was also the only SRG with a Repbase hit using Censor, also to a Gypsy LTR retrotransposon. These results indicate that YIL092W, a *Saccharomyces*-conserved SRG, is likely derived from a Gag TE gene.

### Profile-profile alignments identify homologs of YCL021W-A and YLR036C outside Saccharomyces suggestive of transposable element origin

We aimed to identify direct evidence of genus-conserved SRGs originating from transposition events. Towards this end, we searched for homologs of each genus-conserved SRG for which we had not identified extrageneric homologs in any prior analysis against homology groups constructed from all ORFs of each sequenced yeast genome in Opulente at al. 2024^21^, using profile-profile alignment with hh-suite. As a negative control, we also aligned all homology group profiles against profiles constructed from the reverse sequence of each tested SRG. Among negative controls, no alignments had e-values below 10^−6^; we therefore set this as the threshold for follow-up among the candidate SRGs.

Two SRGs, YCL021W-A and YLR036C had alignments passing this threshold. Both genes are within 1 kb of a transposable element in the S. cerevisiae genome and thus likely exist in a transposition hotspot. The SRG YCL021W-A matched to a homology group in *Zygosaccharomyces* (e-value 7 × 10^−7^) while YLR036C matched to a group in *Naumovozyma* (e-value 8.9 × 10^=8^). For both homology groups matching to an SRG, we used the genes within the group as queries in a BLASTP search (e-value threshold 10^−4^) against the full set of genomes assembled in Opulente et al. 2024 to attempt to identify additional potential homologs. For YCL021W-A, this search identified matches only in the *Zygosaccharomyces*, *Torulaspora* and *Zygotorulaspora* genera (Supplemental Figure 7), all in single copy except for two copies in *Zygosaccharomyces sapae*. For YLR036C, this search identified no matches outside the three sequenced species in the *Naumovozyma* genus, at single copy in *N. dairenensis* and *N. baii* and tandem duplicate in *N. castellii* (Supplemental Figure 7). The limited phylogentic distribution of both these gene families is consistent with a scenario of two independent horizontal transfer events into post-WGD genomes.

We examined the vicinity of the potential homologs of YCL021W-A and YLR036C to assess whether they too were located in transposable element hotspots. All matches to YCL021W-A outside *Saccharomyces* were located between homologs of the *S. cerevisiae* genes YPR120C and YPR122W. This region contains three Ty1 LTRs in the SGD reference genome and is upstream of a cysteine tRNA; regions upstream of tRNA genes tend to be Ty insertion hotspots.^52^ According to YGOB^27^, many budding yeast genera contain a cysteine tRNA in this sytenic block including *Zygosaccharomyces* and *Torulaspora,* the genera in which the YCL021W-A matches are found. In contrast, the matches to YLR036C are located between homologs of *S. cerevisiae* genes YOR025W and YOR027W, which does not appear to be a transposition hotspot. This syntenic block is not near a tRNA in any species on YGOB, not does it have any transposable element features annotated on SGD. However, YLR036C does have homologs in transposable element hotspots within *S. cerevisae* itself. YLR036C has four annotated paralogs in the *S. cerevisiae* reference genome, all of which lack BLASTP homologs outside *Saccharomyces* (YIL089W, YPL257W, YPR071W, YIL029C). Though present on four different chromosomes, all five paralogs are located in likely transposable element hotspots: YLR036C, YPL257W and YIL029C are adjacent to Ty1 LTRs; YIL089W is adjacent to a Ty4 LTR; YPR071W is not adjacent to any annotated transposable element feature but is immediately upstream of a serine tRNA. This genomic pattern suggests that YLR036C either has transposition capacity or was descended from a gene with transposition capacity. Yet, YLR036C synteny is conserved through the genus, with a YLR036C ortholog present between YLR035C and YLR037C in all *Saccharomyces* species. Conservation within a single locus suggests that YLR036C provides a benefit for the organism and is not merely a selfish transposable element gene. In any case, we observe for both YCL021W-A and YLR036C multiple associations with transposable element hotspots, a strong hh-clust match with a narrow taxon in addition to *Saccharomyces*, and no identifiable homologs outside these taxa; these observations favor the hypothesis of an origin by transposable element mediated horizontal transfer rather than *de novo* birth within *Saccharomyces*.

## Competing Interest Statement

A.-R.C. is a member of the Scientific Advisory Board for Flagship Labs 69, Inc. (ProFound Therapeutics).

## Acknowledgments

This work was supported by funds provided by the National Science Foundation grant MCB-2144349 awarded to A.-R.C.

## Supplementary Figures

**Supplementary Figure 1:**
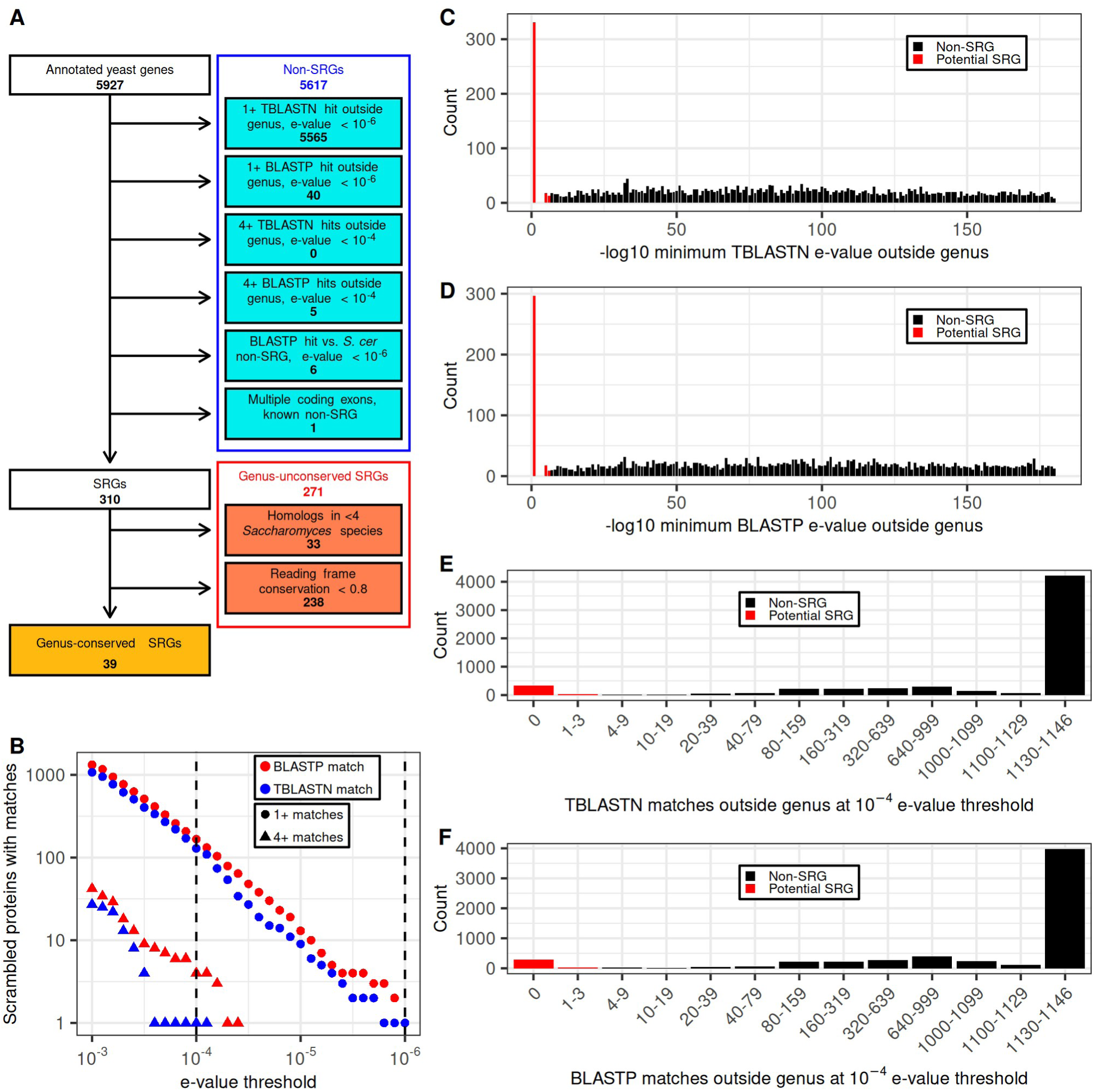
Identifying SRGs within the Saccharomyces genus. A) Flowchart describing how SRGs were identified and separated into genus-conserved and genus-unconserved classes. B) The number of scrambled protein sequences (negative controls, N=5904) with either at least one or at least four BLASTP or TBLASTN matches to budding yeast genomes outside Saccharomyces, out of 1146 possible, across a range of e-value thresholds. Dashed lines indicate the chosen thresholds for inferring a high confidence match, 10^−4^ for four matches and 10^−6^ for a single match; these imply <0.1% false discovery rate. C-D) Distribution of minimum TBLASTN (C) or BLASTP (D) e-values among S. cerevisiae annotated genes (N=5904) against genomes outside Saccharomyces. E-values below 10^−6^ are inferred to indicate high confidence homology, implying that the gene is not a SRG. E-F) Distribution of counts of genomes with TBLASTN (E) or BLASTP (F) matches (e-value < 10^−4^) among annotated S. cerevisiae genes (N=5904) against 1146 budding yeast genomes outside Saccharomyces. Genes with four or more matches are inferred to be non-SRGs.

**Supplementary Figure 2:**
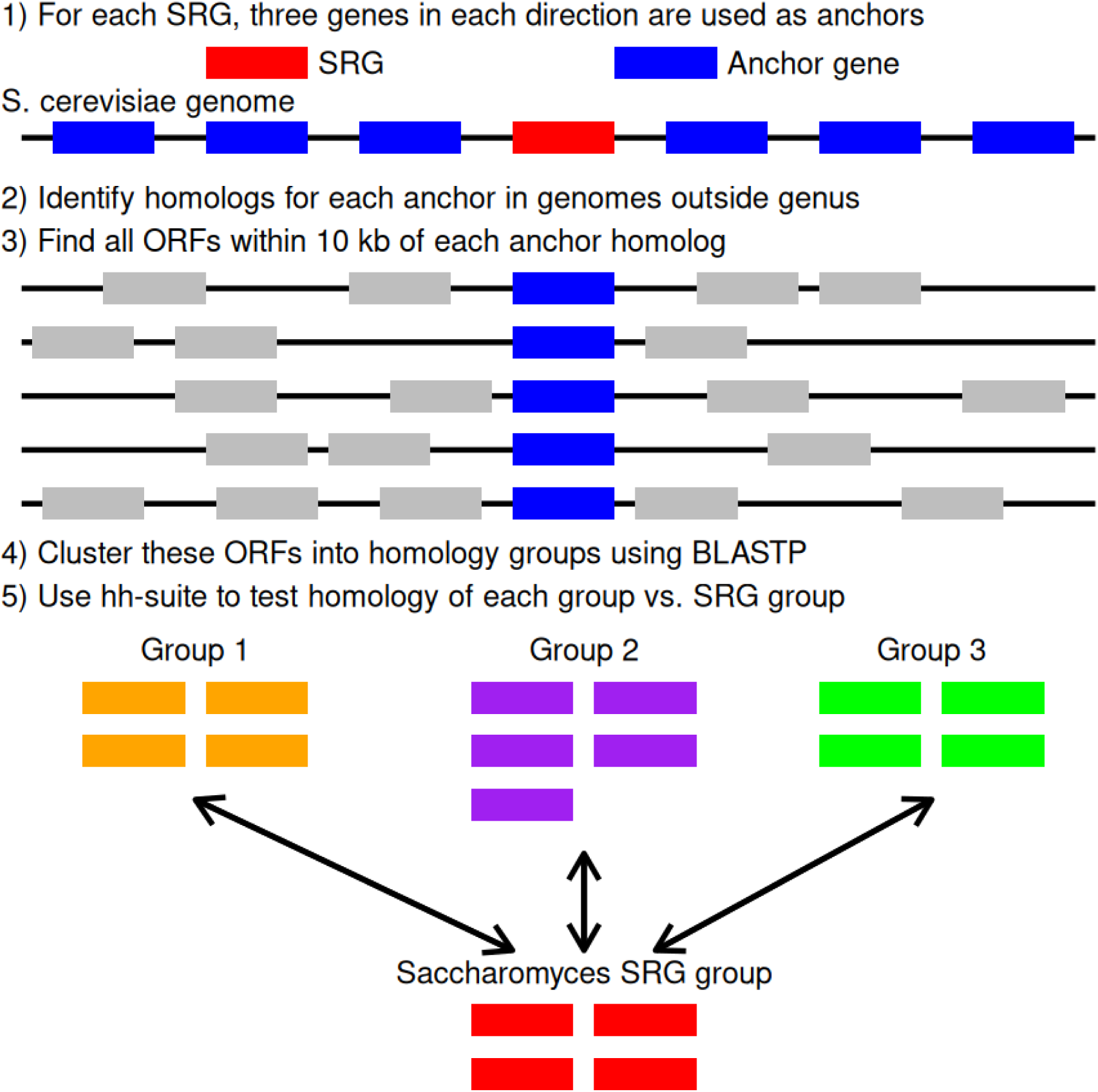
A systematic approach to identify homologs of SRGs in syntenic regions. For each SRG, the three closest annotated genes in each direction are used as potential anchors. In each comparison genome outside the genus, homologs are identified for each potential anchor and all ATG-to-stop ORFs within 10 kb of each anchor homolog are identified. These ORFs are then clustered into homology groups using BLASTP. A homology group for the SRG is constructed from its homologs within the genus. Then, hh-suite is used to attempt to align each extrageneric homology group to the SRG homology group.

**Supplementary Figure 3:**
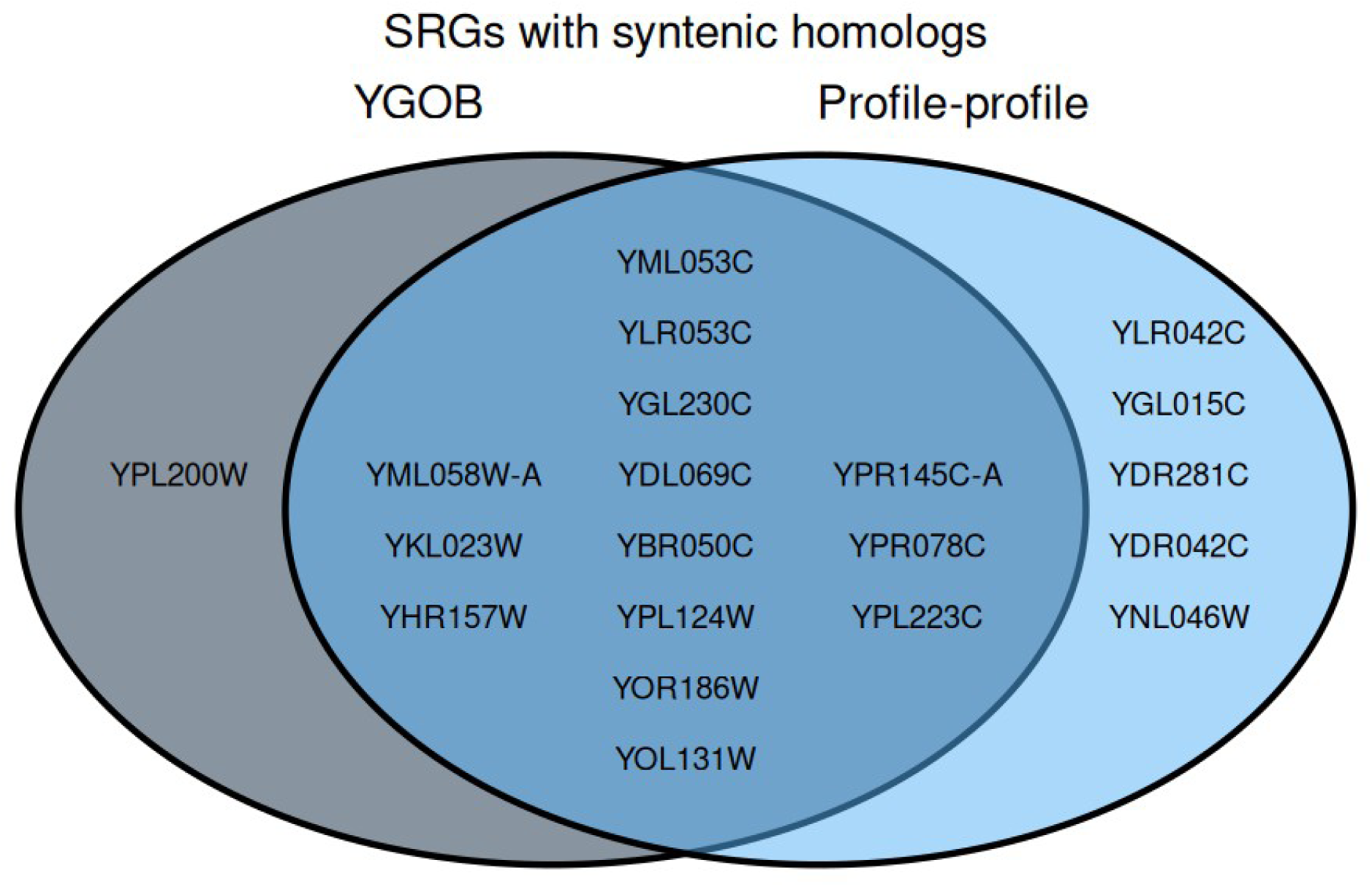
Comparison between automated profile-profile alignment approach and Yeast Genome Order Browser. Among SRGs conserved in Saccharomyces, the genes with syntenic homologs outside Saccharomyces according to YGOB, our systematic profile-profile alignment approach, or both, are listed.

**Supplementary Figure 4:**
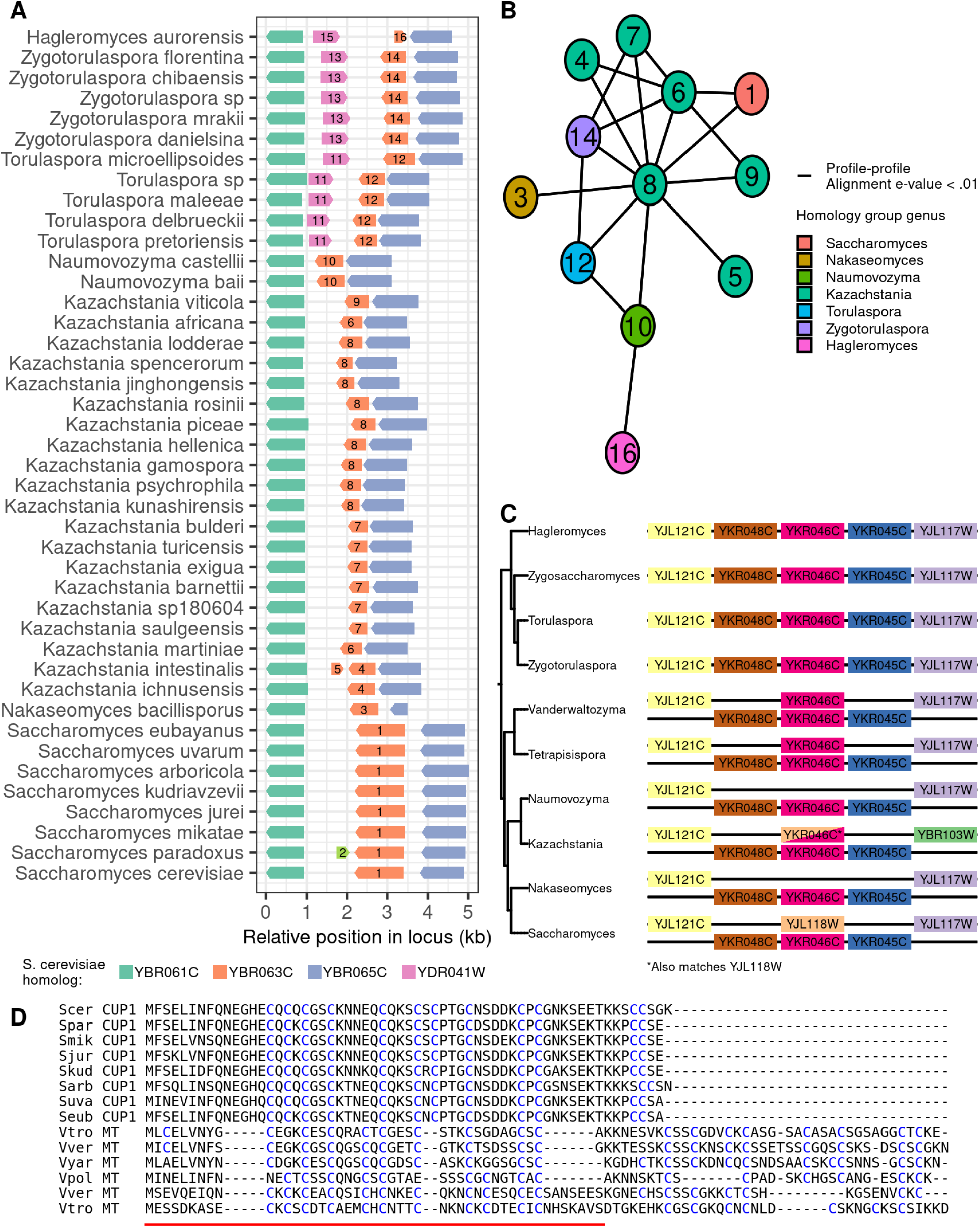
Identification of vertically descended SRGs by close examination of individual cases. A) Syntenic block bounded by homologs of YBR061C and YBR065C in all post-WGD genomes in which at least one ORF longer than 300 nt is present in the block. This block contains the SRG YBR063C in Saccharomyces. All ORFs longer than 300 nt are represented except homologs of YBR062C. Color indicates the closest homolog in S. cerevisiae if there is at least one match with BLASTP e-value < 1 × 10^−4^ in a genome vs. genome search; otherwise, ORFs are colored based on inferred homology by profile-profile alignment as indicated in Supplementary Figure 4B. The number in each ORF indicates the homology group it is assigned to. B) Graph of homology groups found in the same syntenic block as YBR063C where edges are drawn between groups that have pairwise profile-profile alignments with e-value < .01. Homology group ids indicated on each node correspond to Supplementary Figure 4A. C) Phylogenetic pattern of syntenic blocks containing homologs of YJL118W and YKR046C, where gene order within each block is indicated. Each represented block corresponds to the general pattern of gene order among species in the indicated genus. The label for each ORF indicates the closest S. cerevisiae homolog if there is at least one match with BLASTP e-value < 1 × 10^−4^ in a genome vs. genome search; otherwise, ORFs are labeled if they have a profile-profile alignment match with e-value <.01 to an S. cerevisiae gene in the same syntenic block. D) Multiple alignment between Saccharomyces CUP1 and metallothioneins in Vanderwaltozyma genus identified by synteny with known N. glabrata metallothionein MT1. The red line indicates the extent of the local profile-profile alignment given by hh-suite; the multiple alignment follows this profile-profile alignment in this region, while the remainder of the MSA follows a multiple alignment generated using MUSCLE. Note that some Vanderwaltozyma species have multiple copies of the gene at the locus.

**Supplementary Figure 5:**
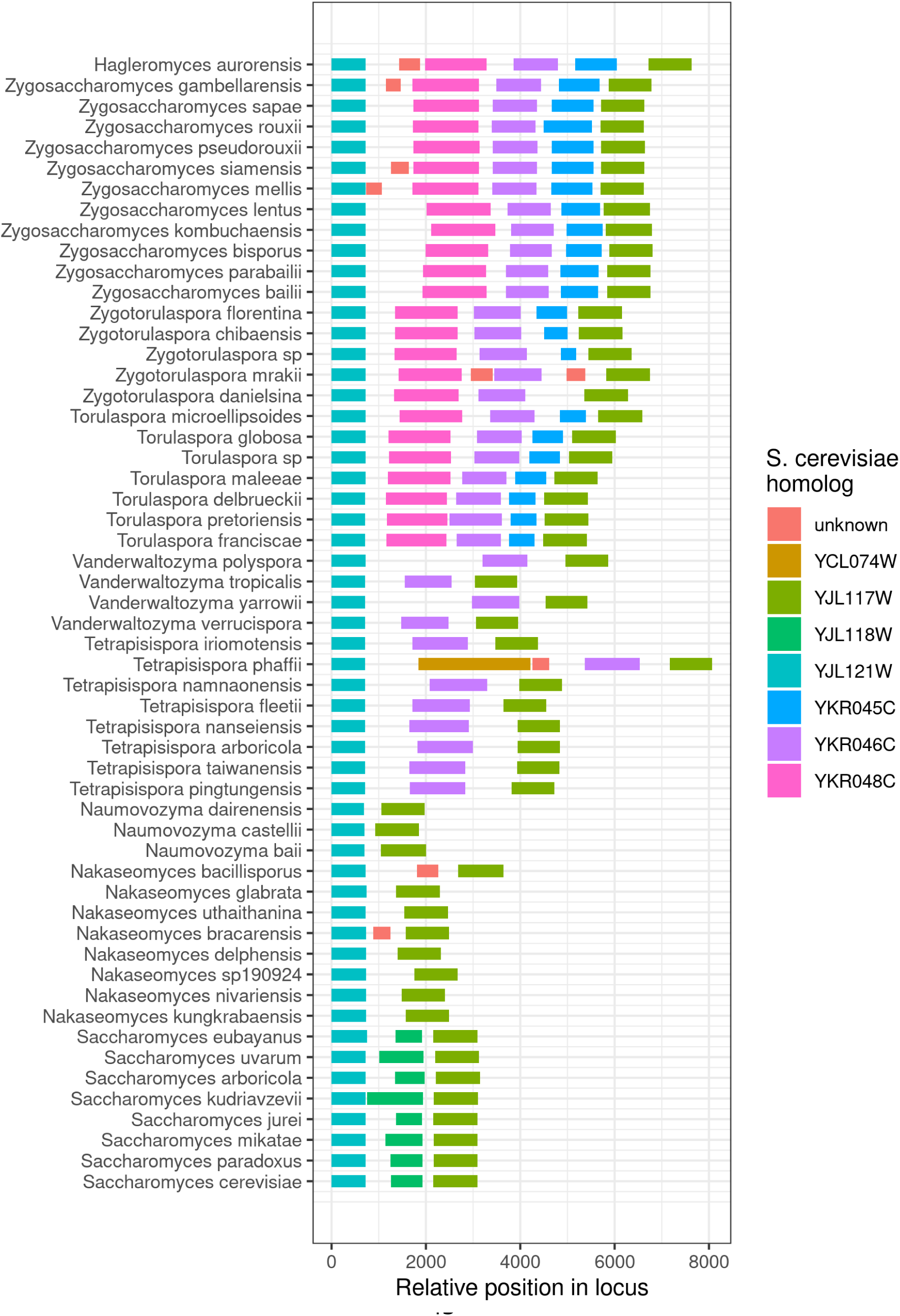
The syntenic block between YJL117W and YJL121W is conserved in most post-WGD species. Syntenic block bounded by homologs of YJL117W and YJL121W in all post-WGD genomes in which this block exists. The SRG YJL118W exists within this block in Saccharomyces. All ORFs longer than 300 nt are shown. Color indicates the closest homolog in S. cerevisiae if there is at least one match with BLASTP e-value < 1 × 10^−4^ in a genome vs. genome search.

**Supplemental Figure 6:**
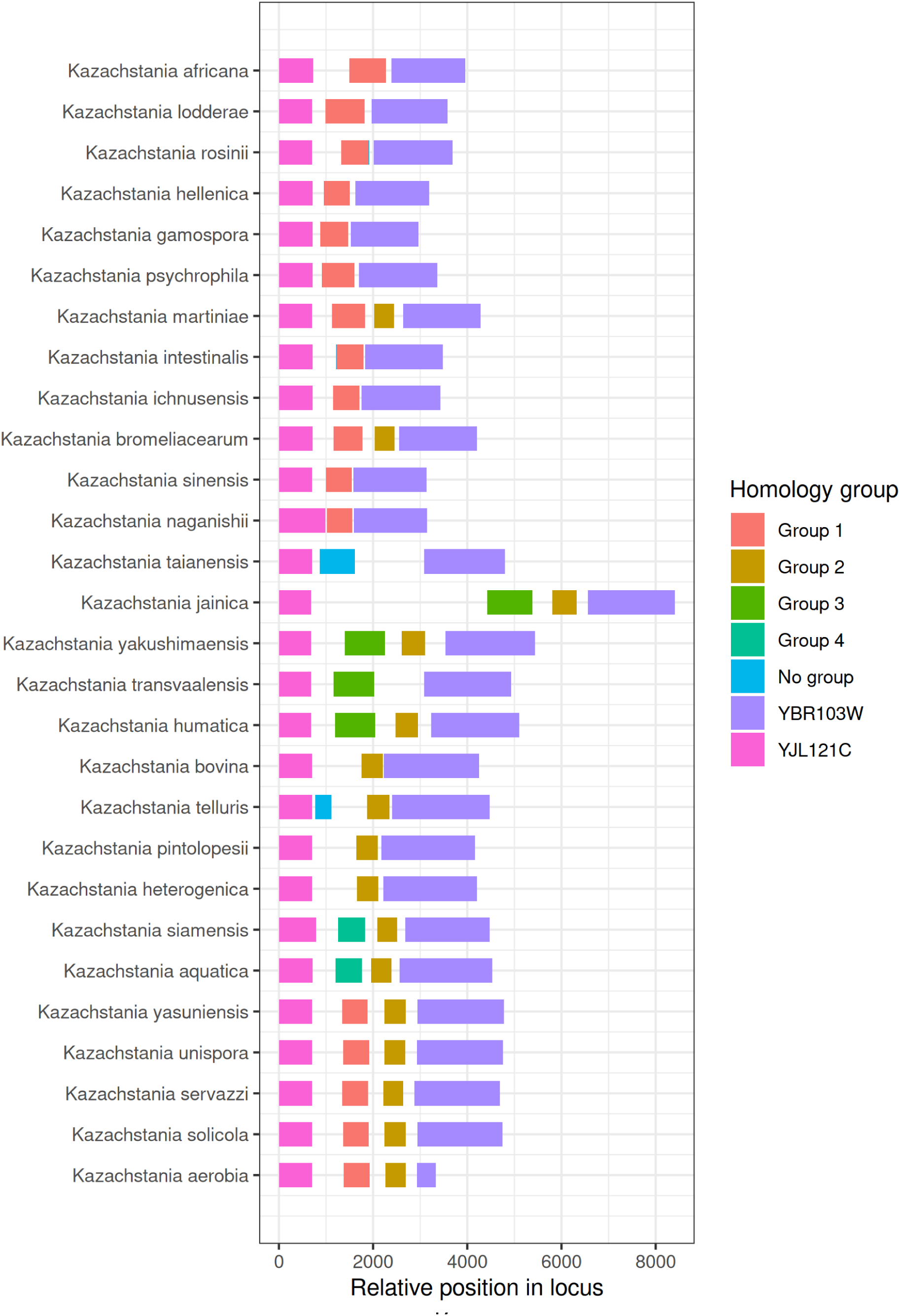
In Kazachstania species, YJL121C is adjacent to a YBR103W homolog rather than YJL117W. Syntenic block bounded by homologs of YJL121W and YBR103W in Kazachstania species. All ORFs longer than 300 nt are shown. Color indicates the homology group each ORF within the block is assigned to.

**Supplemental Figure 7:**
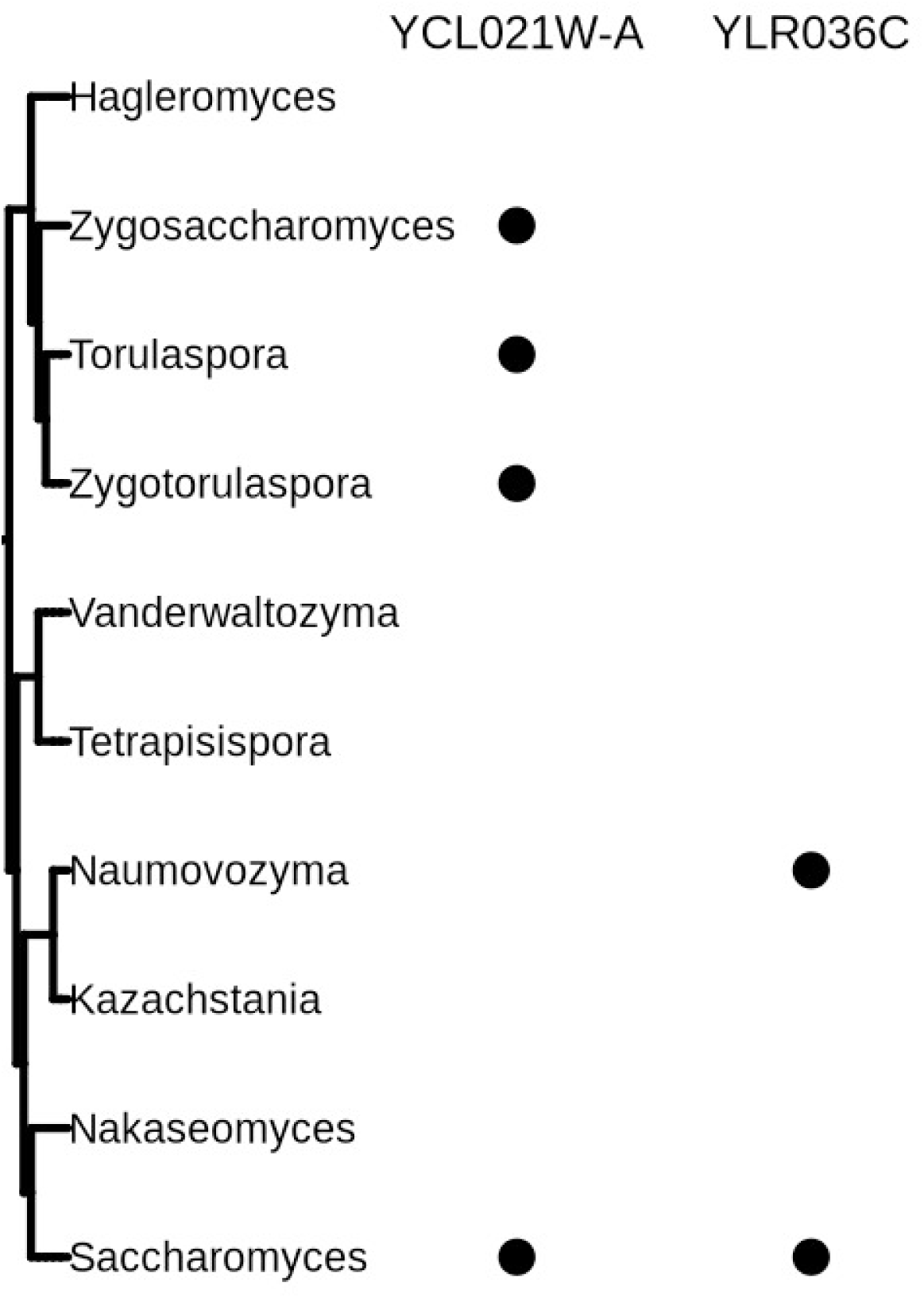
YLR036C and YCL021W-A are SRGs with likely transpsoosable element associated origins. Genera with identified homologs for YCL021W-A and YLR036C. A genera is considered to have an identified homolog if it has a homology group with profile-profile e-value < 10^−6^ in hhsuite against the Saccharomyces homology group of the given gene, or if it has any ORF with a BLASTP match e-value < 10^−4^ with any of the ORFs in the matching (non-Saccharomyces) homology group.

**Supplemental Figure 8:**
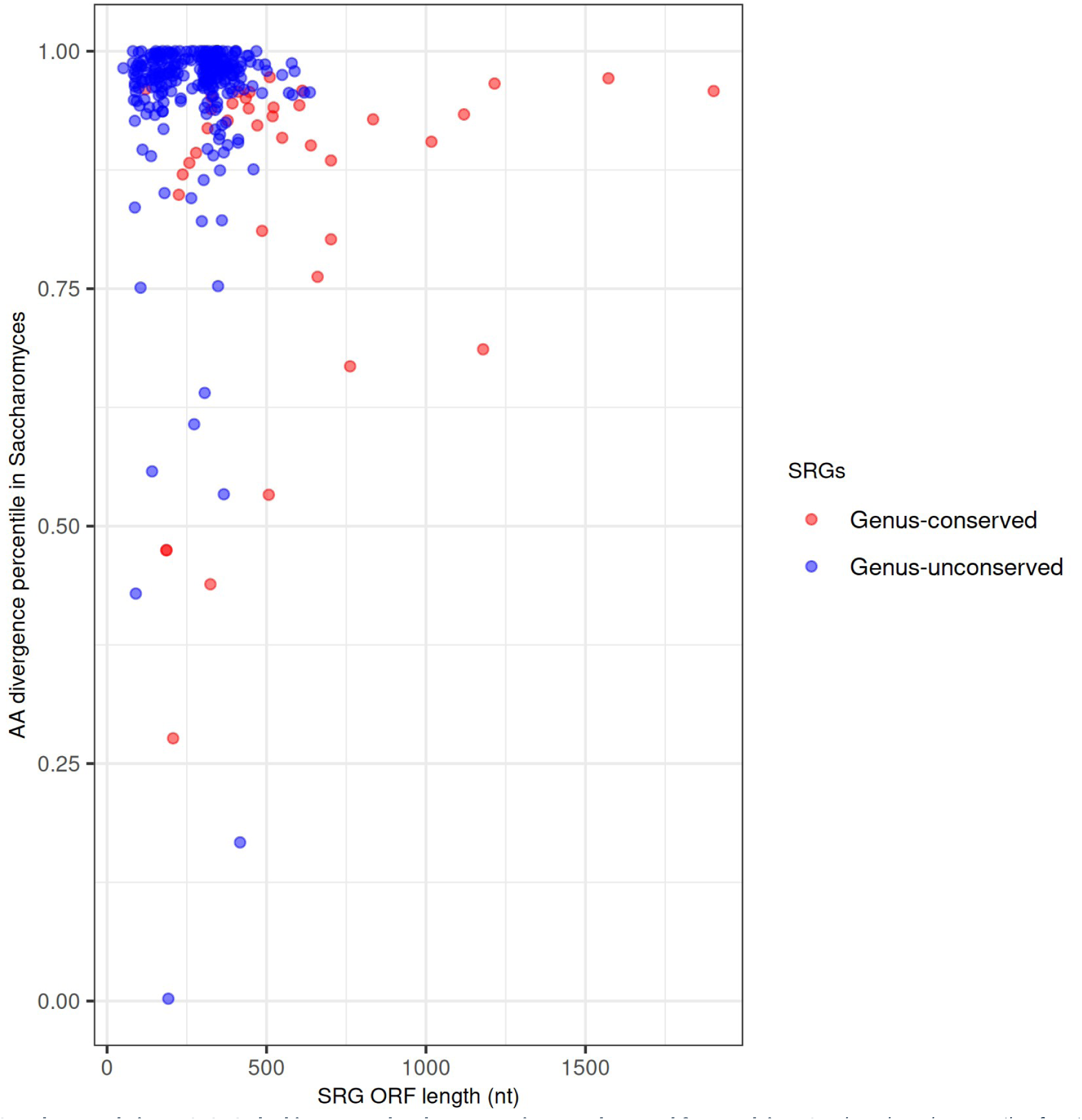
SRGs lacking genus-level conservation are short and fast evolving. ORF length and percentile of amino acid divergence within Saccharomyces among annotated genes for all genus-conserved (N=39) and genus-unconserved (N=271) SRGs.

